# ADPKD variants in the PKD2 pore helix cause structural collapse of the gate and distinct forms of channel dysfunction

**DOI:** 10.1101/2024.09.12.612744

**Authors:** Orhi Estarte Palomero, Paul G. DeCaen

## Abstract

PKD2 is a member of the polycystin subfamily of transient receptor potential (TRP) ion channel subunits which traffic and function in primary cilia organelle membranes. Millions of individuals carry pathogenic genetic variants in PKD2 that cause a life-threatening condition called autosomal dominant polycystic kidney disease (ADPKD). Although ADPKD is a common monogenetic disorder, there is no drug cure or available therapeutics which address the underlying channel dysregulation. Furthermore, the structural and mechanistic impact of most disease-causing variants are uncharacterized. Using direct cilia electrophysiology, cryogenic electron microscopy (cryo-EM), and super resolution imaging, we have discovered mechanistic differences in channel dysregulation caused by three germline missense variants located in PKD2’s pore helix 1. Variant C632R reduces protein thermal stability, resulting in impaired channel assembly and abolishes primary cilia trafficking. In contrast, variants F629S and R638C retain native cilia trafficking, but exhibit gating defects. Resolved cryo-EM structures (2.7-3.2Å) of the variants indicate loss of critical pore helix interactions and precipitate allosteric collapse of the channels inner gate. Results demonstrate how ADPKD-causing these mutations have divergent and ranging impacts on PKD2 function, despite their shared structural proximity. These unexpected findings underscore the need for mechanistic characterization of polycystin variants, which may guide rational drug development of ADPKD therapeutics.

**Regarding polycystin nomenclature:** The revised and current IUPHAR/BPS nomenclature creates ambiguity regarding the genetic identity of the polycystin family members of transient receptor potential ion channels (TRPP), especially when cross-referencing manuscripts that describe subunits using the former system^1^. Traditionally, the products of polycystin genes (e.g., PKD2) are referred to as polycystin proteins (e.g., polycystin-2). For simplicity and to prevent confusion, we will refer to the polycystin gene name rather than differentiating gene and protein with separate names— a nomenclature we have recently outlined (Annual Reviews in Physiology, Esarte Palomero et al. 2023)^2^

## INTRODUCTION

Renal polycystins (PKD1, PKD2) are ion channel subunits with associated genetic variants that cause autosomal dominant polycystic kidney disease (ADPKD)^3,4^. ADPKD is lethal monogenetic disorder characterized by progressive cyst development that precipitates renal failure^5,6^. More than 12 million individuals carry disease causing variants in PKD1 or PKD2, which accounts for the majority (>95%) clinically reported cased of ADPKD^5^. Although ADPKD is commonly considered a loss-of-function disease putatively driven by haploinsufficiency or defective protein expression— the mechanistic impact of most polycystin variants remain uncharacterized^6,7^. There is no cure for ADPKD, and current treatments focus on managing symptoms and indirect means of slowing disease progression^8,9^. Thus defining the structural and functional impact of disease-causing polycystin variants is critical to inform the development of targeted therapies.

ADPKD is considered a “channelopathy” and a “ciliopathy”— meeting both criteria for a disease that is caused by ion channel and cilia dysregulation^10,11^. Renal polycystins traffic to the primary cilium of collecting duct principal cells, where they form homomeric (PKD2) and heteromeric channel complexes (PKD1-PKD2)^12-14^. While the characteristics of native heteromeric PKD1-PKD2 channel conductance are poorly understood, recent work suggests that the complex is not constitutively active, but rather activated after cleavage sequence of the PKD1 N-terminus^12^. In contrast, homomeric PKD2 channels are voltage-gated and Ca^2+^-modulated when assessed directly from the primary cilia membrane using submicron diameter electrodes^13-15^. Primary cilia are small (d ∼ 200 nm, l ∼ 2-8 μm), singular projections that are highly insulated from the cell body^16^. PKD2 contributions to plasma membrane current is nominal when assessed natively in kidney principal cells or when the channel is overexpressed^13^. While the high-resolution cryogenic electron microscopy (cryo-EM) structures of homomeric polycystins have provided a molecular bases for the present study, our understanding of the structural motifs responsible for channel function and assembly is limited^17-20^. Given their tenability to structural analysis using cryo-EM and functional analysis within the primary cilia, we will primarily consider the impact of PKD2 variants within the context of homomeric channels.

Previously, we used direct cilia electrophysiology, and cryo-EM variant structure determination to define how variants in the PKD2 TOP domain causes structural destabilization and gating defects^21^. In this manuscript, we apply these techniques along with super resolution cilia trafficking analysis to evaluate PKD2 pathogenic (https://pkdb.mayo.edu/variants) germline missense variants (F629S, C632R, R638C) found within pore helix 1 (PH1) of the channels pore domain (**Supplemental Figure 1**)^22^. Our results define mechanistic differences among these loss-of-function variants, despite impacting the same structural motif within the PKD2 channel. Alterations in the variant cryo-EM structures and their biophysical gating defects implicate a long range allosteric mechanism controlling the opening of the channel pore. Significance of the findings underscores the need to further characterize the diversity in polycystin variant molecular dysfunction to guide rational approaches ADPKD drug development.

## RESULTS

### Destabilizing effects of PH1 variants on protein stability, channel assembly and structure

To biochemically assess the impact of ADPKD missense variants, we transiently expressed these channels in HEK cells and isolated PKD2 protein using DDM (n-dodecyl β-d-maltoside) detergent. The oligomeric assembly of WT, F629S and R638C protein samples is apparent by their monodispersed elution fractions as evaluated by size-exclusion chromatography (SEC), whereas C632R protein was prone monomeric disassociation (**Figure 1A)**. This variant-specific destabilizing effect was supported by dye-based thermal denaturing temperature gradients (**Figure 1B**). Here, the C632R channel was highly unstable at physiological temperatures (37-38C°), whereas the remaining variants only denatured after heating the samples beyond 50C°. After plunge freezing variant channels reconstituted C8 amphipols and collecting negative staining data sets, the unfolded C632R protein was apparent from the disordered densities observed in micrographs, whereas assembled channel particles were readily visualized from the other variants samples. These characteristic precluded C632R variant samples from further structural characterization and we proceeded to single particle cryo-EM analysis of the F629S and R638C variants. Between 437K-665K particles of the variant channels were collected, processed, and refined to resolve their molecular structures at 3.2Å (F629S) and 2.7Å (R638C) overall resolution (**Figure 1C, Supplementary Figure 2A, B**). Each channel subunit encodes six transmembrane spanning helices (S1-S6) which fold into structural domains: voltage sensor (VSD, S1-S4), tetragonal opening for polycystins (TOP), pore (PD, S5-S6) and cilioplasmic (N- and C-termini). Like previously resolved WT channel structure, the PKD2 variant structures forms a domain-swapped tetrameric channel with N- and C-termini (**Figure 1D**)^17-20^. The VSD of the F629S and R638C variants structures are captured in the deactivated state with S4 gating charges (K572, K575) oriented for hydrogen bonding and cation-π interactions that facilitate gating charge transfer (**Supplementary Figure 3A, B**). The deactivated VSDs are expected given that polycystins are voltage-gated channels that open at positive voltages and the structures were capture without a membrane potential^13,14,23,24^. The ion conducting pathway within the PD of homomeric polycystin channels have two restriction sites which were proposed to function as “gates” (**Figure 2A-C**)^19,25^. The putative external gate is found within the ion selectivity filter (L641-N643) that is scaffold by two reentrant pore helices (PH1, PH2), and the internal gate is located at tetrameric S6 junction formed hydrophobic residues (L677-N681). Both gates of F629S and R638C variant channels are captured in non-conducting closed states based their PDs minimum radii (R_min_=0.55-0.62Å) that are too narrow for partially hydrated cations (Na^+^, K^+^, Ca^2+^) to permeate (**Figure 2B, C**, (**Supplementary Figure 4A-C**). As expected, several local PH1 molecular interactions are absent from the F629S and R638C variant structures compared to the previously resolved WT PKD2 channel (**Figure 2A**)^19^. On the lipid-facing side of the PH1 in the F629S channel structure, the large phenyl sidechain density which is normally found buried in a hydrophobic cleft formed by cilia membrane and non-polar residues (L609, A612) of the S5 was replaced by an unfavorable polar serine hydroxyl. In the R638C structure, the guanidium functional group is replaced by sulfhydryl, disrupting a hydrogen-bond tripartite network formed with carboxylate and hydroxyl sidechains of the PH1 (E631, T635), and selectivity filter (D643) normally found in WT channels. Despite the localized disruptive these interactions, the pitch and secondary fold of the PH1 helix are preserved in the F629S and R638C variant structures, and only caused a small dilation (ΔR_min_<0.24Å) in the external gate (**Figure 2B, C**). However, examination of the variant channel’s internal gates reveals key differences in L677 and N681 sidechains rotamers that effectively double the length its restriction (6-8Å) along the S6 (**Figure 2B, C**). Here, changes in the orientation N681 carboxamides optimizes hydrogen bonding distances (3.4-2.8Å) between each protomer. These interactions stabilize the occluded ion conducting pathway observed in the R636C structure, and also create asymmetry observed in at the inner gate of the F629C channel (**Supplementary Figure 3C**). Based on these biochemical and structural results— ranging from complete channel assembly failure (C632R) to allosteric collapse of internal gate (F629S and R638C)— we investigated the PH1 variant impacts on channel trafficking and biophysical properties in primary cilia membranes.

**Figure 1.**
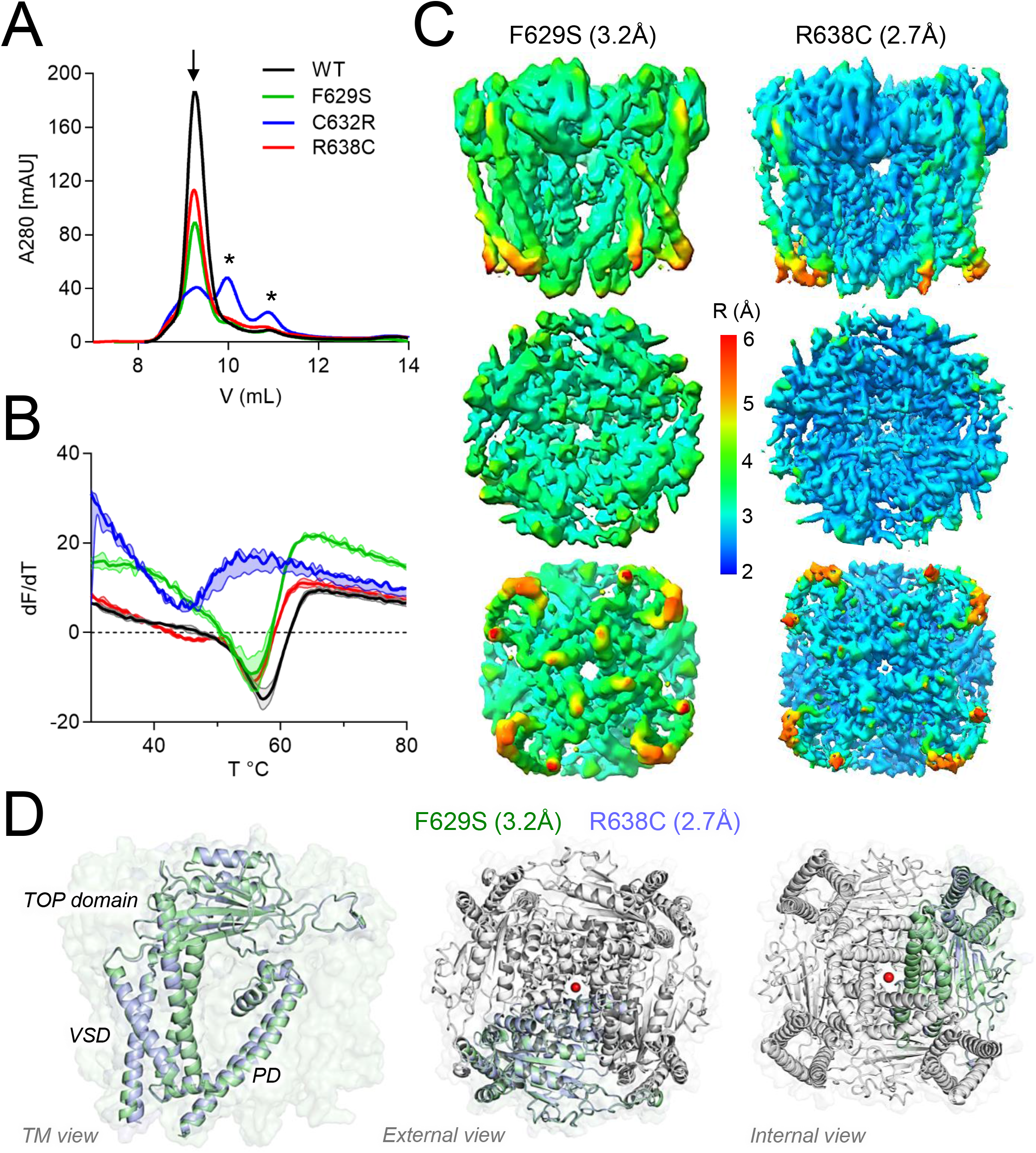
Cryo-EM structural determination of ADPKD-causing PKD2 missense variants and their impact on channel assembly. **A**) Size exclusion chromatography results of WT and PH1 variant channels purified from HEK cells. Peaks indicate the relative abundance of each channel as oligomers (arrow) or protomers (asterisk). **B**) GlowMelt (Biotinum) thermal stability measurement of polycystin protein unfolding. First derivative (slope) of the fluorescence curve (dF/dT) for each channel protein is plotted as a function of temperature (T). Error bars = S.D., N=3 replicates for each channel type. **C**) Global resolution maps of PKD2 F629S and R638C structures determined by cryo-EM. Note, pore domain resolution is between 2.8-3.1Å and 2-2.4Å for the F629S and R638C structures, respectively. **D**) Overall structure of the variant channels. *Left*, transmembrane view and structural alignment of the F629S and R638C channels, highlighting the domains within a single channel subunit. *Right*, external and internal views of the channels demonstrating the domain swapped assembly of the channel.

**Figure 2.**
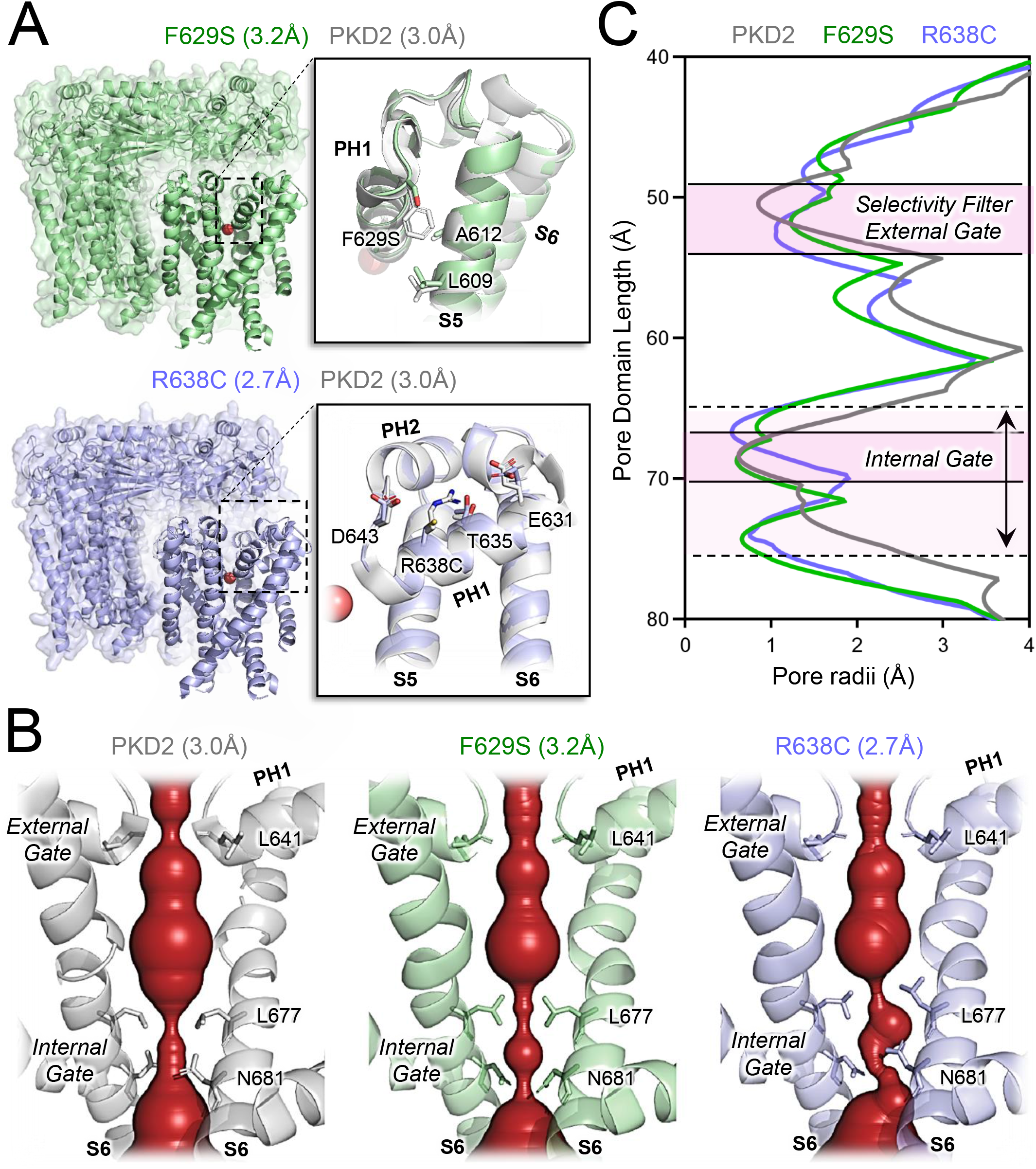
Allosteric structural effect of PH1 variants on the PKD2 ion conducting pathway. **A**) Structural alignment of the variant and WT channels (PDB: 5T4D, Cao et al 2016). Inset, expanded views the pore domains showing the location of the ADPKD-associated variants highlighting the missing pore helix chemical interactions. **B**) HOLE analysis (Smart et al 1996) of the PKD2 pore domains highlighting the changes in the internal and external gates. **C**) Analysis of the polycystic pore radii plotted along their length. Note the elongated pore restriction at the internal gates (double pointed arrow) of the F629S and R638C compared to the WT channels.

### PH1 missense variants cause divergent effects on primary cilia trafficking and channel gating

Using super resolution 3D-structured illumination microscopy (3D-SIM), we compared primary cilia trafficking of the PKD2 pore helix variants. PKD1 and PKD2 null HEK cells stably expressing ciliary ARL13B-GFP to visualize primary cilia were transiently transfected with HA-tagged variant channel construct prior to fixation, immunolabeling and colocalization analysis (**Figure 3A, B**)^21^. As expected, misfolded and unassembled C632R channel subunits variants do not localize to the primary cilia and remain sequestered in the cell. In contrast, the structured and assembled F629S and R638C variants traffic to the ciliary organelle membrane, like WT channels (**Figure 3A, B**). Overexpression of the PH1 variants significantly impaired primary cilia length of these cells, where the C632R variant had the greatest impact (WT L_cilia_ = 4.7μm, C632R L_cilia_ = 2.4μm). To assess the impact of the variants on channel function, we used microelectrodes to form high resistance seals with the primary cilia membrane and performed single channel voltage clamp recordings (**Figure 3C-E**)^13^. Voltage-dependent channel openings were evident from cilia expressing F629S and R638C channels (**Figure 3C**). In agreement from the cilia trafficking defects observed by super resolution, no channel opening events were detected from cilia patch clamp recordings from cells expressing the C632R variant. Voltage dependence of opening (V_1/2_) of the F629S and R638C channels were positively shifted (ΔV_1/2_ = 27-32mV) compared to WT, which increased the free energy of gating (ΔGº) by +1.2 kcal/mol, +1.8 kcal/mol, respectively (**Figure 3D, Table 1**). The unitary conductance (γ) of both variants were significantly decreased, suggesting small reduction in rate of cation permeation though the pore. Taken together, the super resolution and electrophysiology data sets establish divergent loss of channel mechanisms caused ADPKD variant. While the F629S and R638C variant results in a partial loss of function, producing significant energy barrier to opening the PKD2 channel, the C632R variant causes a complete loss of cilia localization through its failure to traffic of this organelle.

**Table 1.**
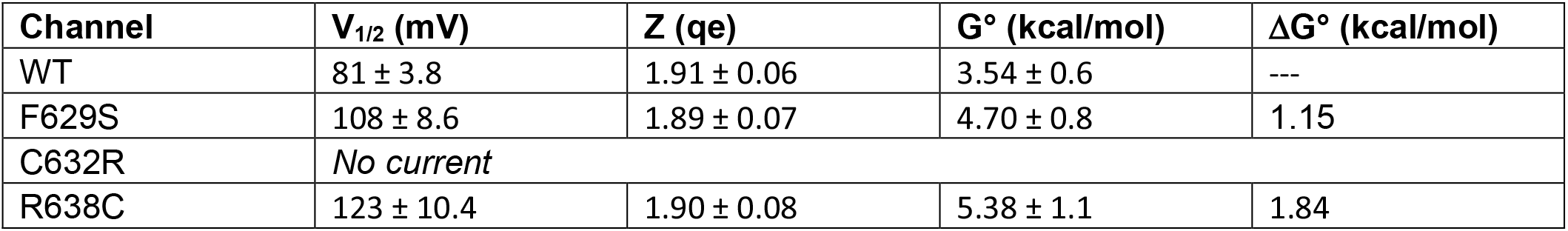
Gating properties of PKD2 ADPKD PH1 variants. Boltzmann parameters (V_1/2_) and (Z) resulting from fitting the voltage-dependent opening relationship of PKD2 channels reported in Figure 3. G° indicates free energy of channel opening quantified by the Gibbs free energy equation described in the methods section. ΔG° indicates the change in free energy induced by the variant. Error = S.D.

| Channel | $V_{1/2}$ (mV) | $Z$ (qe) | $G^\circ$ (kcal/mol) | $\Delta G^\circ$ (kcal/mol) |
| --- | --- | --- | --- | --- |
| WT | $81 \pm 3.8$ | $1.91 \pm 0.06$ | $3.54 \pm 0.6$ | --- |
| F629S | $108 \pm 8.6$ | $1.89 \pm 0.07$ | $4.70 \pm 0.8$ | 1.15 |
| C632R | *No current* |  |  |  |
| R638C | $123 \pm 10.4$ | $1.90 \pm 0.08$ | $5.38 \pm 1.1$ | 1.84 |

**Figure 3.**
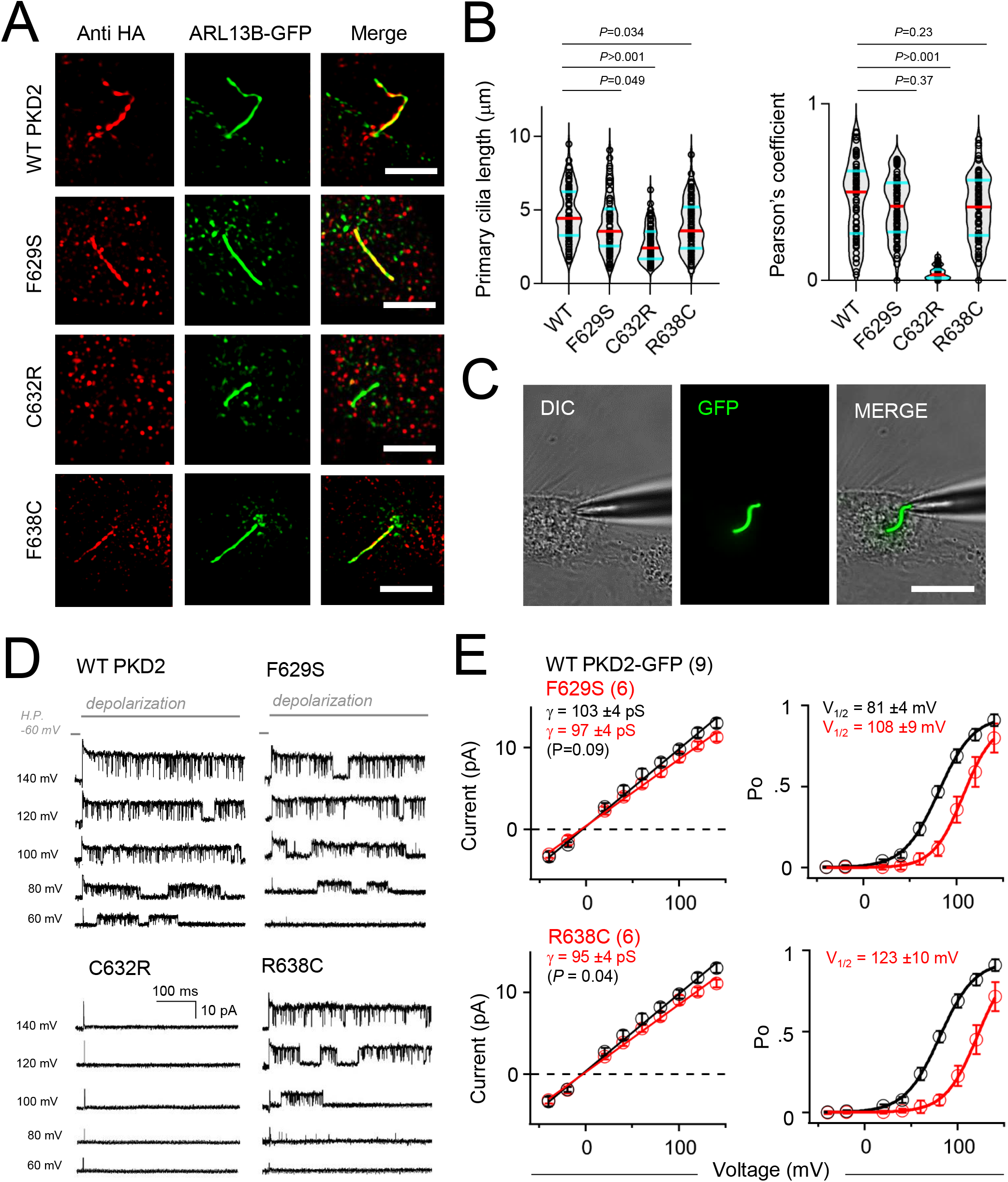
ADPKD-causing PKD2 missense variants cause partial to complete loss of channel function through impaired trafficking defects or impaired gating. **A**) Super resolution SIM (structured illumination microscopy, Scale bar = 3 μm) images of HEK PKD1^Null^:PKD2^Null^ cells stably expressing ARL13B-GFP (primary cilia reporter) and transiently transfected HA-PKD2 channels immunolabeled with anti-HA antibody (red). **B**) Primary cilia length and fluorescence colocalization analysis using the Pearson’s correlation coefficient. **C**) Image a voltage clamped HEK cell primary cilium illuminated under fluorescence. **D**) Example PKD2 WT and variant single channel records recorded from the cilia and activated by the indicated depolarization steps. **E**) *Left*, single channel current magnitudes comparing WT and variant channels. Unitary conductance was estimated by the slope (γ) of the linear fit. *Right*, Open probability (Po) plotted as a function of voltage and fit to a Boltzmann equation to estimate the slope (Z) and half-activation voltage (V_1/2_). Error bars = S.D. and the number of cilia records (N) for each data set is indicated within the parenthesis. *P*-values indicate results from two tailed, unpaired Student’s t-tests comparing WT and variants channel data sets.

## DISCUSSION

### Summary of findings

Using multidisciplinary methods which include super resolution imaging, single-particle cryo-EM structural determination and cilia microelectrode electrophysiology— we have defined the molecular mechanism and functional consequences of three PKD2 missense variants that cause ADPKD. Even though all three mutations are located within the same PH1 structural motif, they have divergent mechanistic impacts. Through biochemical and 3D-SIM analysis, we determine that C632R variant causes a complete loss of function through destabilized oligomeric assembly which precipitates channel exclusion from the primary cilia. In contrast, the F629S and R638C variants cause partial loss of function— having normal trafficking and assembly, but impaired voltage-dependent gating. This assessment is supported by features observed in our determined F629S and R638C cryo-EM structures. These missense variants break a critical PH1 interactions but do not cause catastrophic changes in the fold and assembly of the channel pore. Rather, the F629S and R638C variants cause an allosteric structural change to pore lining S6 at the internal gate— doubling the length its hydrophobic septum. The unitary conductance of these variants also decreased contributing an additional loss of function effect. Discussed below are the significance of these finding related to PKD2 channel molecular regulation and developing polycystin variant-targeted drugs for the ADPKD population.

### Implication for polycystin structural gating mechanisms

The molecular mechanisms controlling opening of the polycystin pore domain are poorly understood. As identified in the initial cryo-EM structural determination of PKD2, the pore domain’s ion-conducting pathway are constricted at internal and external sites, which are proposed to serve as gates^19^. While it is widely accepted that the external site is the ion selectivity filter, its putative function as a gate is controversial^25,26^. The pore helices scaffold the selectivity filter/external gate, but we did not find significant changes at this site in our pore helix 1 variant structures. While we reported a small dilation (ΔRmin<0.24Å) at this site, our interpretation is limited by the resolution limits of the cryo-EM data sets. PH1 variants caused a reduction unitary conductance, a functional observation consistent with selectivity filter defects limiting cation flow through the pore. In contrast to the external gate, the variants caused significant changes (lengthening, asymmetry) to the internal gate formed by the crossing of S6 helices. These inner gate structural defects were functional validated by voltage dependent gating defects recorded from the variant channels at the cilia membrane. The role of pore helices as structural regulators of channel opening is further supported by results from an unbiased mutagenic screen of the related PKD2L1 pore domain^27^. Taken together, these results are most consistent with of a “one-gate” mechanism— where long-range coupling energetics between the pore helices are transferred to the inner gate within the pore. *How can variants within the PH1 alter the inner gate’s function, given the sites are separated by long molecular distances (≈20Å)*? The upper and lower gates are likely allosterically coupled via an interaction network that involves residues at the selectivity filter (F641) and pore helix 1 (R638) of one subunit and pore helix 2 (F646) and the pore-lining S6 helix (F661 and V665) of a neighboring subunit (**Supplementary Figure 3B**). The ADPKD R638C variant removes a cation-π interaction between R638 and F646 and reduced the coupling between the pore helices of two adjacent subunits. Whereas the polarity of the F629S serine side chain unfavorably dips into a hydrophobic pocket formed by the membrane and non-polar S6 residues of the same subunit. We hypothesize that this mutation alters the conformational coupling between the VSD and PD, stabilizes the non-conductive state of the pore observed in our F629S cryo-EM structure. Importantly, both variants impact this gating mechanism through discreate chemical interactions, without altering structure of the local pore helix, or the overall quaternary channel assembly. Taken together, our study of the PH1 variants has elucidated the unexpected biophysics governing channel gating, conductivity and provided a mechanistic explanation for their loss of function that drives ADPKD progression.

### Impact on the rationale for polycystin-targeted ADPKD drug development

There is currently no drug cure for ADPKD, but development of pharmaceuticals which activate variant polycystins or reinstating their channel function may present a viable therapeutic strategy. This view is supported by in vivo studies in which de-repression and rescue of polycystin gene expression ameliorates cyst growth in mouse models of ADPKD^28,29^. These results indicate that either PKD1 or PKD2 reactivation through genetic or pharmaceutical intervention is sufficiently attenuates disease progression^28,29^. We have determined the PKD2 variants F629S and R638C traffic to the primary cilia but have defective gating properties and reduced ion conductivity. Based on our findings, activating these functional and cilia-localized polycystins using prototypic channel modulators may be a viable drug strategy. Alternatively, reinstating PKD2 R636C trafficking through the design of molecular chaperones (“correctors”) that stabilize the channel protein may also prove beneficial. A proof of concept of VX-407, a prototypic drug under clinical trials, corrects defective PKD1 folding to treat ADPKD in patients with a subset of PKD1 variants^30^. However, the efficacy of this drug against the ADPKD population with PKD2 variants is unreported. Further elucidation of mechanistic differences amongst disease-causing variants and the polycystin pharmacophore are needed to evaluate prototypic pharmaceutics for the genetically diverse ADPKD patient cohort. Alternatively, developing patient genotype-specific (i.e., variant-targeted) therapeutics may inform a personalized medicine approach as drug development strategy for ADPKD.

## METHODS

### Expression, purification, and denaturing of PKD2 channels

PKD2 WT and point mutant variants (F629S, C632R, R638C) were expressed by transient transfection in Expi293F GnTl^-/-^ cells using an N-terminal Strep-tagged MBP-TEV-PKD2 construct (hPKD2: 52-793) under control of a CMV + β-globin intron promoter. Expi293F GnTl^-/-^ cells were grown at 37 °C to 3×10^6^ cells/mL density and transfected with PEIMax (DNA:PEIMax 1:5). 20-22h post transfection sodium butyrate was added to boost protein expression (10 mM). Cells were harvested 72 h post transfection, washed with PBS, and stored at -80 °C until further use. Cell pellets were resuspended in Buffer A (25 mM HEPES pH = 7.4, 150 mM NaCl, 1 mM TCEP, 1 mM CaCl_2_, 10% glycerol) supplemented with complete protease inhibitors. The cells were homogenized by sonication at 30% power and supplemented with n-dodecyl-β-D-maltoside (DDM) in Buffer A to a final concentration of 2% w/v. Membranes were solubilized by gentle nutation at 4 °C for 2 h and cell debris was removed by centrifugation at 20000 g for 30 minutes at 4 °C. The cleared lysate was applied to 0.5 ml of equilibrated Strep-Tactin Superflow affinity resin by gravity flow. The column was washed with 30 CV of Buffer A + 0.05% DDM/0.005% CHS and eluted in 5 CV of Buffer A supplemented with 5 mM D-desthiobiotin. Protein concentration was estimated by A280 followed by supplementation with amphipol A8-35 (1:3 w/w) and incubation for 1 h at 4 °C. Detergent was removed with two batches of SM2 Biobeads (10 mg/0.1 mL) by gentle nutation at 4 °C, first for 1 h and then overnight. The MBP solubility tag was cleaved simultaneously with the last batch of Biobeads by the addition of TEV Protease. PKD2 52-793 reconstituted in A8-35 was polished by size exclusion chromatography in a Superdex 200 10/300 GL column using Buffer B (25 mM pH = 7.4, 150 mM NaCl, 1 mM TCEP, 1 mM CaCl_2_). Peak fractions eluting at ∼9.5 mL post injection was pooled, concentrated with 100 kDa cutoff centrifugation filters and used for CryoEM sample preparation. Thermal shift experiments were performed in a Thermo Fisher QuantStudio 7 qPCR instrument at 2 °C/min temperature ramp from 20 °C to 95 °C using MicroAmp Fast 96-well PCR plates. 10 μg of SEC purified PKD2 variants in A8-35 were mixed to a final concentration of 1X GloMelt dye (33021-T) and 50 nM ROX dye internal standard in 20 μL of buffer (25 mM HEPES pH =7.4, 150 mM NaCl and 1 mM CaCl_2_). Experiments were performed in technical triplicates.

### Cryo-EM Sample Preparation and Data Collection

Samples were plunge frozen in a Leica EM GP2 at 5 °C and >95% humidity and -183 °C liquid ethane temperature. PKD2 (53-792) variants reconstituted in amphipol A8-35 (0.5-1.0 mg/mL) were applied to a glow discharged (air, 15 mA, 20 s) Quantifoil Holey Carbon 2/1 copper 300 mesh grids. 3.5 μL of sample was applied to the carbon side and the grid was immediately blotted for 2.5-3.5 s at 42 mm/2 mm blot distance. Preliminary grid screening and data collection were conducted at the Pacific Northwest Cryo-EM Center (PNNC) using SerialEM software. Datasets were collected on a Titan Krios G3i equipped with a K3 detector and energy filter 10 eV slit. Movie stacks composed of 50 subframes and 60 e^-^/Å total dose were collected at 130000x magnification on super resolution mode (0.3235 Å pixel size). The target defocus acquisition range was set between -0.6 and -2.4 μm.

### Cryo-EM Data Processing

Data processing was performed in CryoSPARC, including the initial preprocessing motion correction, dose weighing and CTF estimation and micrograph curation steps.

PKD2 R638C Variant: 16586 movies were collected. 750 particles from 40 micrographs were picked manually to create initial templates. These templates were used to auto pick particles from 1000 micrographs and create new templates. The second generation of templates was used to auto pick the full 14292 curated set of micrographs. 2084635 particles were extracted with an 864-pixel box size, down-sampled to 432 pixels and subjected to two rounds of 2D classification (50 classes). The top 12 classes (665417 particles) were selected and used to create an ab-initio volume. Homogenous refinement imposing C4 symmetry yielded 2.7 Å GSFSC resolution reconstruction.

PKD2 F629S Variant: 9145 movies were collected from replicate grids. The curated 8508 micrograph stack was used for blob picking polycystin particles (130-170 Å diameter). 2660324 picks were extracted with a 864-pixel box size downsized to 432 px and subjected to two rounds of 2D classification (50 classes). The top 13 classes (408506 particles) were selected and used to create an ab-initio volume. Homogenous refinement imposing C1 and C4 symmetry yielded 4.6 Å and 3.2 Å resolution reconstruction, respectively. B-factor sharpening 160.2 applied to C4 symmetric map.

### Model Building and Structure Validation

Alphafold2 models (PKD2 residues 180-925) containing the variant of interest were pruned in PHENIX^31^ to remove low confidence residues. The resulting PKD2 variant models, comprised of residues E213-K292 and N303-Q694, was manually fitted into the Cryo-EM map using ChimeraX^32^. Model refinement and validation was performed using ISOLDE^33^ which led to the additional pruning of the TM2-TM3 linker region (Residues E494-F503). The TM2-TM3 linker resides on an area of the map with >5 Å local resolution were not adequately modeled (SI Local Resolution Maps).

### Generation of HEK PKD1^Null^:PKD2^Null^ cells lines stably expressing the ARL13B-GFP cilia label

Using our previously generated CRISPR/Cas9 gene edited HEK PKD2^null^ cell line, we introduced nonsense mutations to both PKD1 alleles to create a PKD1^null^:PKD2^nul^l cell line. HEK 293 PKD2^null^ cells were electro-transfected with PKD1 sgRNAs (caccGCATAGGTGTGGTTGGCAGC and aaacGCTGCCAACCACACCTATGC) with the All-in-one Cas9 plasmid (Addgene). Cells generated from single cell clones were selected after 4 weeks of expansion under puromycin selection in a 96-wells plate. A HEK PKD1^Null^:PKD2^Null^ clone was verified by extracting the genomic DNA and sequencing for the introduced STOP codons within PKD1 and PKD2 genes using the following primers: PKD1 fwd (CTGATGGCTTAGGCCCCTACTG); PKD1 rev (CCTGGGTCTCGGTAGATGAACG); PKD2 fwd (AGCCTCAGGGCACAGAACAG); and PKD2 rev (CCACACTGCCCTTCATTGGC). Plasmids for mammalian expression of ADP Ribosylation Factor Like GTPase 13B C-terminally tagged with GFP (ARL13B-GFP) were custom synthesized (Vector Builder) and subcloned into third-generation lentiviral packaging plasmids used for stable expression contained: pMDLg/pRRE (Addgene), reverse transcriptase pRSV-Rev (Addgene), and envelope expressing plasmid pMD2.G (Addgene). LentiX-293T cells (Takara) were transfected with polyethylineimine (Polysciences) at a 4:1:1:1 ratio of the transgene and viral packaging constructs. Supernatants were collected 48 and 72 hours post transfection and filtered through a 0.45 μm syringe filter. Lentiviral supernatant was concentrated 100 times using 1 volume of PEG-it (System Biosciences) virus precipitation solution and 4 volume of lentivirus-containing supernatant. The PEG-it and supernatant mixture were kept at 4 degrees for 24 hours and centrifuged at 1500 rpm for 30 minutes. The pellet containing lentivirus was resuspended with 1/100th volume of PBS of the original supernatant volume. Cells were infected with the lentivirus supernatant; ARL13B-GFP expression was selected using culture media containing puromycin (2 μg/ml) for 30 to 90 days. Cells were then fluorescence-activated cell sorted (BD FacsMelody) at 5000 to 10,000 counts per minute to enrich for the transgene expression. Stable cell lines were cultured in Dulbecco’s modified essential medium (DMEM) supplemented with 10% fetal bovine serum (FBS) and 100 units/ml penicillin, 100 units/ml streptomycin and 1 μg/ml puromycin selection antibiotic. Expression plasmids encoding N-terminal HA tagged (hemagglutinin, YPYDVPDYA) human PKD2 were created using the Gibson assembly method. Missense variants were generated using standard, site-directed mutagenesis. Cells were transfected transiently transfected (Lipofectamine 2000, Invitrogen) 24-48 hours prior to electrophysiology and super resolution experiments.

### Structure Illumination Microscopy

PKD1^Null^:PKD2^Null^ cells lines stably expressing the ARL13B-GFP cilia label and transiently transfected with HA-PKD2 channels were fixed with 40% paraformaldehyde (PFA), permeabilized with 0.2% Triton X-100, and blocked by 10% bovine serum albumin in PBS. Cells and tissue were mounted on glass slides and treated with Fluoshield from Sigma-Aldrich (St. Louis MO). Primary cilia localization of the variant channels within were visualized by [1/1000] treated with anti-HA (rabbit) antibodies conjugated with DyLight™ 680 (600-444-384, Rockland). Super resolution images were obtained using an inverted N-SIM Nikon microscope configured for in the Deep 3D-SIM imaging with a 60x silicon oil immersion, 1.3 N.A. objective. Super resolution images using the SIM method were captured under 100× magnification with piezo stepping. Confocal images were further processed with FIJI ImageJ (National Institutes of Health) and Imaris 9.3 (Oxford Instruments).

### Electrophysiology recordings of polycystins from primary cilia membranes

All research chemicals used to for our electrophysiology experiments were purchased and supplied Millipore-Sigma. Voltage clamp experiments capturing transiently transfected PKD2 whole cell currents were made from PKD1^Null^:PKD2^Null^ cells lines stably expressing the ARL13B-GFP cilia label. Cells were seeded onto glass coverslips and placed in a perfusion chamber for voltage clamp recordings in the on-cilia configuration^16^. Ciliary PKD2 single channel events were recorded using borosilicate glass electrodes polished to resistances of 14–22 MΩ. The pipette solution contained (in mM): 90 NaMES, 10 NaCl, 10 HEPES, 10 Na4-BAPTA (Glycine, N, N’-[1,2-ethanediylbis(oxy-2,1-phenylene)]bis[N-(carboxymethyl)]-,tetrasodium); pH was adjusted to 7.3 using NaOH. The bath solution contained 120 KCl, 20 NaCl, 10 HEPES, 1.8 CaCl_2_; pH 7.4 with NaOH to neutralize the resting membrane potential. All solutions were osmotically balanced to 295 (±6) mOsm with mannitol. Data were collected using an Axopatch 200B patch clamp amplifier, Digidata 1440A, and pClamp 10 software. Currents were digitized at 25 kHz and low pass filtered at 10 kHz. Cilia membranes were held at -60 mV and activated by 1.5 s voltage steps mV from -40 mV to 160 mV at successive +20 mV steps. Data were analyzed by Igor Pro 7.00 (Wavemetrics, Lake Oswego, OR). Open probability-voltage relationships were fit using the previously described Boltzmann equation. Cilia unitary conductance (γ) were estimated by fitting the single channel current to a linear equation f(x) = γ(Vm)+b. The amount of free energy (G°) required to open WT and variant channels was calculated using this equation: G°=z(F)V_1/2_ where, F is Faraday’s number, z is the estimated charge based slope derived from the constant.

### Statistical Analysis

Statistical methods used to determine significance are described in corresponding figure legends and reported as *P* values. Briefly, electrophysiology and imaging datasets were analyzed (GraphPad or Origen) using one way ANOVA, Student’s t-tests two tailed paired (equal sample sizes) or unpaired (unequal sample sizes).

## Supporting information

PDB Validation Reports

## Acknowledgements

We thank the staff at the Pacific Northwest Cryo-EM facility (PNCC) and Stanford-SLAC Cryo-EM facility (S2C2) for providing training— in particular Marzia Mileto PhD for data collection assistance (Proposal #160398). We thank Jian Payendeh PhD and Thomas Clairfeuillie PhD (Genentech) for their helpful suggestions regarding protein purification methods and single particle analysis strategies. We thank Eduardo Guardarrama for his assistance in analyzing the channel pore structures. The atomic coordinates for the PKD2 F629S and R638C variant channels have been deposited in the protein databank under accension codes PDB 9DLJ (EMDB entry ID EMD-46980) and 9DLI (EMDB entry EMD-46979), respectively.

## FIGURES AND TABLES

**Supplemental Figure 1.**
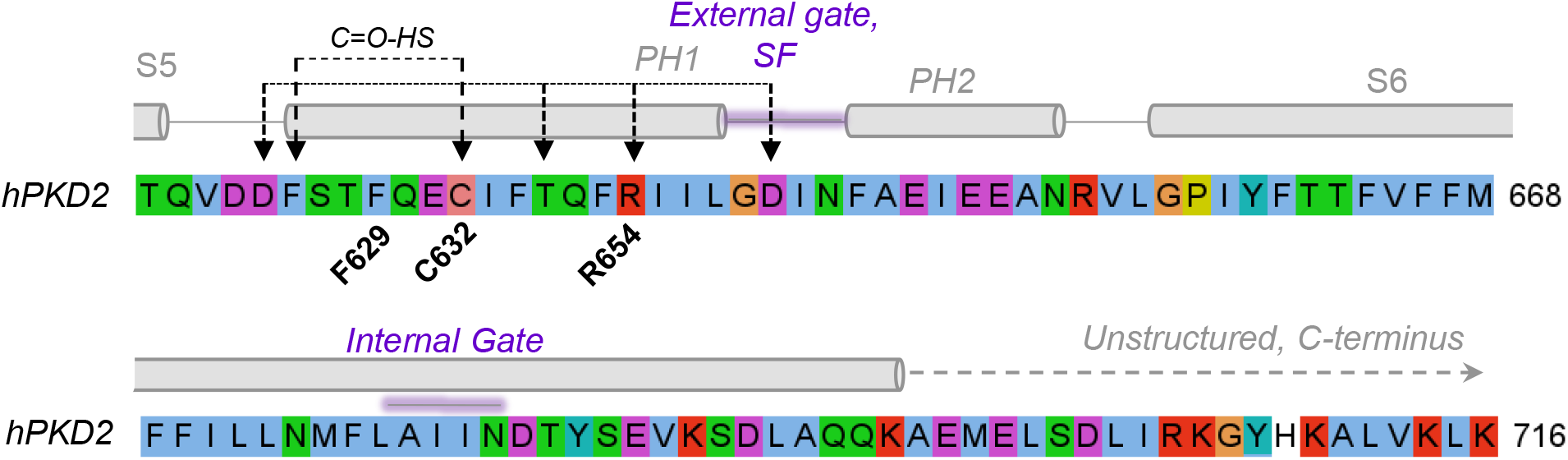
Location of ADPKD-causeing missense variants in the PKD2 pore helix 1. An amino acid sequence rendered in JalView applying the default color scheme: hydrophobic (blue); polar (green); glutamine, glutamate, aspartate (magenta). Special amino acids are designated with their own color: glycine (orange); proline (yellow) and tyrosine or histidine (cyan). The barrels indicate alpha helices found in the PKD2 structure (PDB: 5T4D) and conserved salt-bridge/hydrogen bonds with the external pore domain are indicated by connecting arrows.^34^ Location of PKD2 variants associated with ADPKD are indicated in bold print.

**Supplemental Figure 2.**
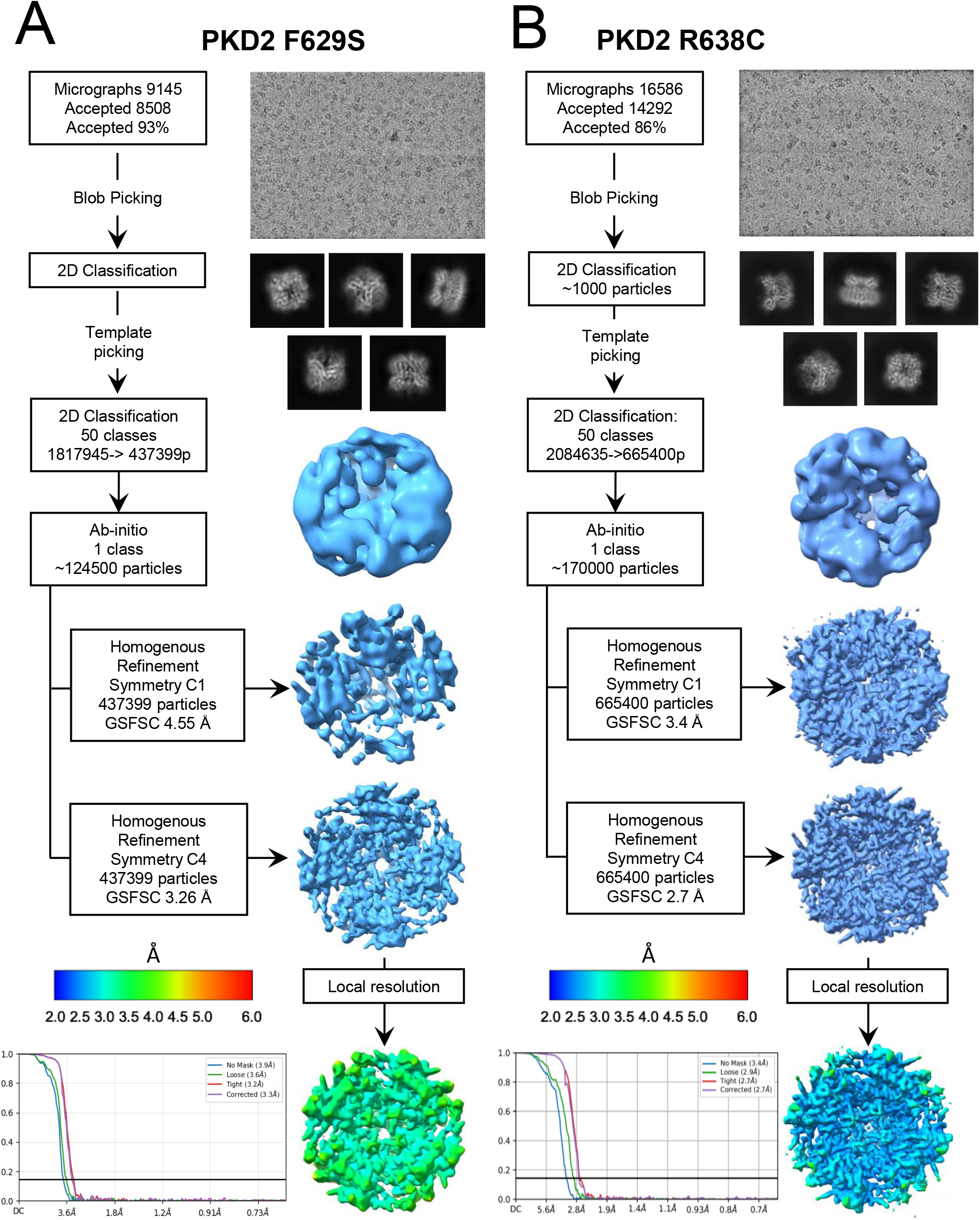
Cryo-EM workflows used to determine the PKD2 F629S and R638C varient structures. Flow charts outlineing the cryo-EM data collection and analysis workflow. Represnetaive micrographs, 2D class averages, and classiications were use to resolve Ab initio class particals. After homogenous refinment, loaclat resolution maps were generated to build and structurel moldles, which were then further refined (See methods section).

**Supplemental Figure 3.**
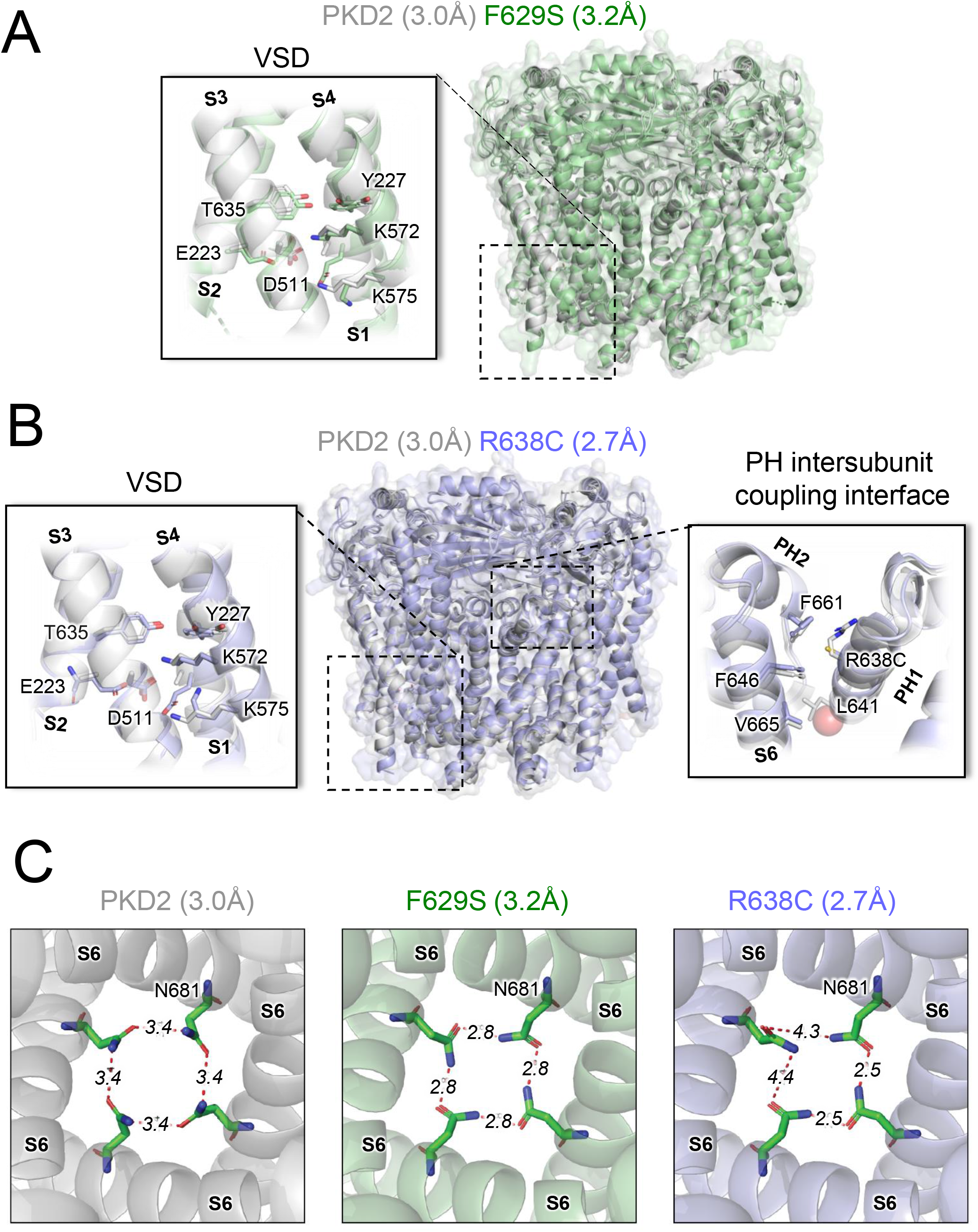
Key structural features of the PKD2 F629S and R638C varient channels. **A, B**) Left, transmembrane views of the voltage sensor domains with the gating charges and interacting hydrogen bond partners highlighted. Note gating charges in the varinat and Wt channel structures are in the deactivated state, below the gating charge transfer center. Right, the pore helix intersubunit coupling interface which is likely affected during opening of the R638C varianst channels. **C**) Intercellular view of the homotypic N681 hydrogen bond arrangement for the WT and varainst structures.

**Supplemental Figure 4.**
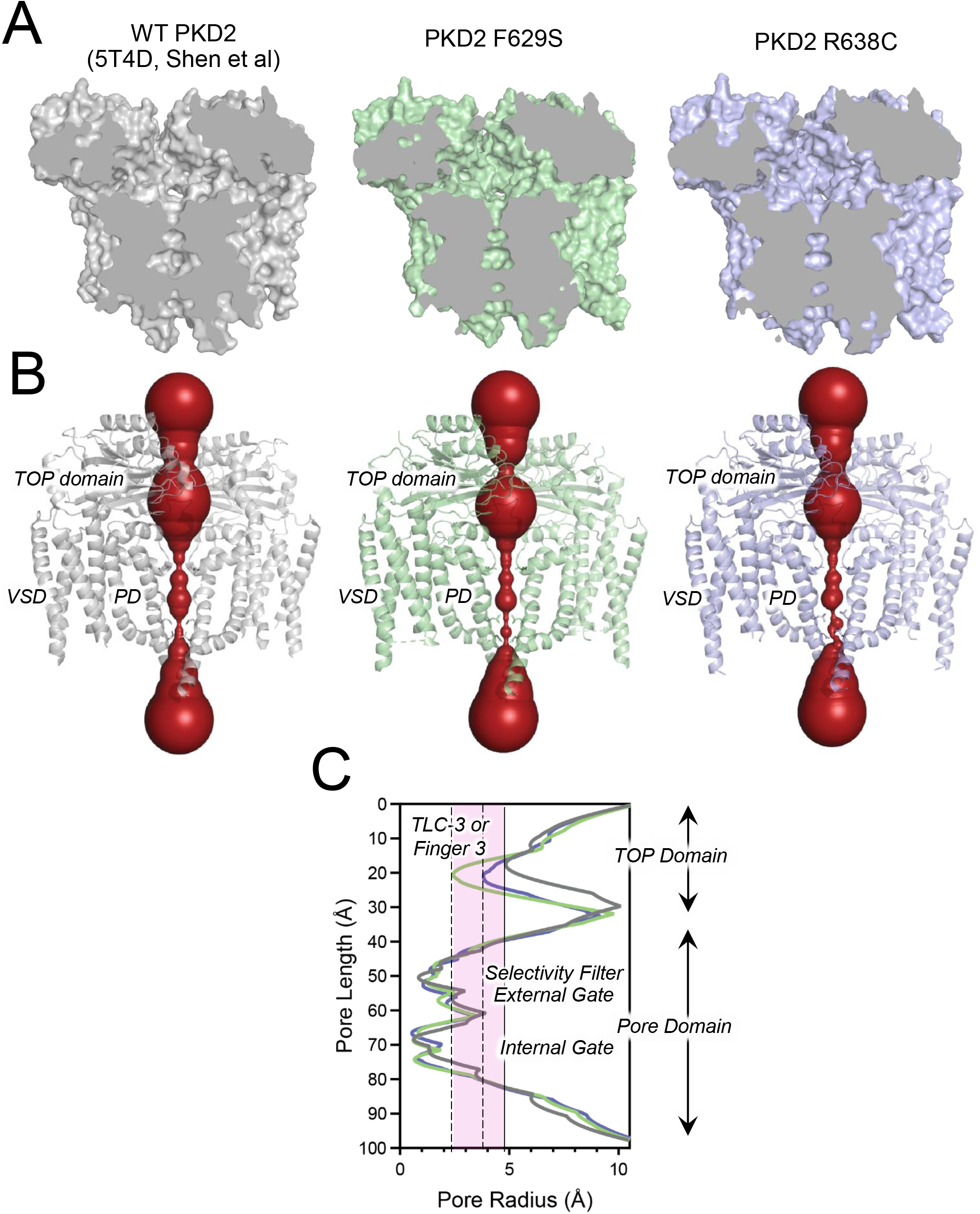
Analysis of the ion conducting pathway in the cryo-EM varaint structures. **A**) Comparision of the ion condcuting pathway from bisected WT and variant polycystin channel structures. **B, C**) Expanding view (TOP domain to pore domain) of the HOLE analysis results (Smart et al 1996) and the pore radius-length plots comparing the WT and variant channels illustrating the changes in the TOP domains.

## REFERENCES

1 Alexander, S. P. et al. THE CONCISE GUIDE TO PHARMACOLOGY 2017/18: Overview. Br J Pharmacol 174 Suppl 1, S1–S16, doi:10.1111/bph.13882 (2017) PMC5650665.

2 Esarte Palomero, O., Larmore, M. & DeCaen, P. G. Polycystin Channel Complexes. Annu Rev Physiol 85, 425–448, doi:10.1146/annurev-physiol-031522-084334 (2023)

3 Hughes, J. et al. The polycystic kidney disease 1 (PKD1) gene encodes a novel protein with multiple cell recognition domains. Nat Genet 10, 151–160, doi:10.1038/ng0695-151 (1995)

4 Mochizuki, T. et al. PKD2, a gene for polycystic kidney disease that encodes an integral membrane protein. Science 272, 1339–1342, doi:10.1126/science.272.5266.1339 (1996)

5 Harris, P. C. & Rossetti, S. Molecular diagnostics for autosomal dominant polycystic kidney disease. Nat Rev Nephrol 6, 197–206, doi:10.1038/nrneph.2010.18 (2010)4050432.

6 Bergmann, C. et al. Polycystic kidney disease. Nat Rev Dis Primers 4, 50, doi:10.1038/s41572-018-0047-y (2018) PMC6592047.

7 Torres, V. E. & Harris, P. C. Polycystic kidney disease: genes, proteins, animal models, disease mechanisms and therapeutic opportunities. J Intern Med 261, 17–31, doi:10.1111/j.1365-2796.2006.01743.x (2007)

8 Chebib, F. T. & Torres, V. E. Autosomal Dominant Polycystic Kidney Disease: Core Curriculum 2016. Am J Kidney Dis 67, 792–810, doi:10.1053/j.ajkd.2015.07.037 (2016)4837006.

9 Chebib, F. T. et al. A Practical Guide for Treatment of Rapidly Progressive ADPKD with Tolvaptan. J Am Soc Nephrol 29, 2458–2470, doi:10.1681/ASN.2018060590 (2018) PMC6171265.

10 Ta, C. M., Vien, T. N., Ng, L. C. T. & DeCaen, P. G. Structure and function of polycystin channels in primary cilia. Cell Signal 72, 109626, doi:10.1016/j.cellsig.2020.109626 (2020) PMC7373203.

11 McConnachie, D. J., Stow, J. L. & Mallett, A. J. Ciliopathies and the Kidney: A Review. Am J Kidney Dis 77, 410–419, doi:10.1053/j.ajkd.2020.08.012 (2021)

12 Ha, K. et al. The heteromeric PC-1/PC-2 polycystin complex is activated by the PC-1 N-terminus. Elife 9, doi:10.7554/eLife.60684 (2020) PMC7728438.

13 Liu, X. et al. Polycystin-2 is an essential ion channel subunit in the primary cilium of the renal collecting duct epithelium. Elife 7, doi:10.7554/eLife.33183 (2018) PMC5812715.

14 Kleene, S. J. & Kleene, N. K. The native TRPP2-dependent channel of murine renal primary cilia. Am J Physiol Renal Physiol 312, F96–F108, doi:10.1152/ajprenal.00272.2016 (2017)5283891.

15 Kleene, N. K. & Kleene, S. J. A method for measuring electrical signals in a primary cilium. Cilia 1, doi:10.1186/2046-2530-1-17 (2012)3539729.

16 DeCaen, P. G., Delling, M., Vien, T. N. & Clapham, D. E. Direct recording and molecular identification of the calcium channel of primary cilia. Nature 504, 315–318, doi:10.1038/nature12832 (2013) PMC4073646.

17 Su, Q. et al. Cryo-EM structure of the polycystic kidney disease-like channel PKD2L1. Nat Commun 9, 1192, doi:10.1038/s41467-018-03606-0 (2018) PMC5864754.

18 Hulse, R. E., Li, Z., Huang, R. K., Zhang, J. & Clapham, D. E. Cryo-EM structure of the polycystin 2-l1 ion channel. Elife 7, doi:10.7554/eLife.36931 (2018) PMC6056229.

19 Shen, P. S. et al. The Structure of the Polycystic Kidney Disease Channel PKD2 in Lipid Nanodiscs. Cell 167, 763–773 e711, doi:10.1016/j.cell.2016.09.048 (2016)

20 Grieben, M. et al. Structure of the polycystic kidney disease TRP channel Polycystin-2 (PC2). Nat Struct Mol Biol 24, 114–122, doi:10.1038/nsmb.3343 (2017)

21 Vien, T. N., Wang, J., Ng, L. C. T., Cao, E. & DeCaen, P. G. Molecular dysregulation of ciliary polycystin-2 channels caused by variants in the TOP domain. Proc Natl Acad Sci U S A, doi:10.1073/pnas.1920777117 (2020)

22 Gout, A. M., Martin, N. C., Brown, A. F. & Ravine, D. PKDB: Polycystic Kidney Disease Mutation Database-a gene variant database for autosomal dominant polycystic kidney disease. Hum Mutat 28, 654–659, doi:10.1002/humu.20474 (2007)

23 Ng, L. C. T., Vien, T. N., Yarov-Yarovoy, V. & DeCaen, P. G. Opening TRPP2 (PKD2L1) requires the transfer of gating charges. Proc Natl Acad Sci U S A 116, 15540–15549, doi:10.1073/pnas.1902917116 (2019) PMC6681712.

24 Vien, T. N., Ta, M. C., Kimura, L. F., Onay, T. & DeCaen, P. G. Primary cilia TRP channel regulates hippocampal excitability. Proc Natl Acad Sci U S A 120, e2219686120, doi:10.1073/pnas.2219686120 (2023) PMC10235993.

25 Cao, E. Structural mechanisms of transient receptor potential ion channels. J Gen Physiol 152, doi:10.1085/jgp.201811998 (2020) PMC7054860.

26 Jara-Oseguera, A., Huffer, K. E. & Swartz, K. J. The ion selectivity filter is not an activation gate in TRPV1-3 channels. Elife 8, doi:10.7554/eLife.51212 (2019) PMC6887487.

27 Ng, L. C. et al. Energetic landscape of polycystin channel gating. EMBO Rep 24, e56783, doi:10.15252/embr.202356783 (2023) PMC10328073.

28 Dong, K. et al. Renal plasticity revealed through reversal of polycystic kidney disease in mice. Nat Genet 53, 1649–1663, doi:10.1038/s41588-021-00946-4 (2021) PMC9278957.

29 Lakhia, R. et al. PKD1 and PKD2 mRNA cis-inhibition drives polycystic kidney disease progression. Nat Commun 13, 4765, doi:10.1038/s41467-022-32543-2 (2022) PMC9376183.

30 Koulen, P. et al. Polycystin-2 is an intracellular calcium release channel. Nat Cell Biol 4, 191–197, doi:10.1038/ncb754 (2002)

31 Liebschner, D. et al. Macromolecular structure determination using X-rays, neutrons and electrons: recent developments in Phenix. Acta Crystallogr D Struct Biol 75, 861–877, doi:10.1107/S2059798319011471 (2019) PMC6778852.

32 Meng, E. C. et al. UCSF ChimeraX: Tools for structure building and analysis. Protein Sci 32, e4792, doi:10.1002/pro.4792 (2023) PMC10588335.

33 Prisant, M. G., Williams, C. J., Chen, V. B., Richardson, J. S. & Richardson, D. C. New tools in MolProbity validation: CaBLAM for CryoEM backbone, UnDowser to rethink “waters,” and NGL Viewer to recapture online 3D graphics. Protein Sci 29, 315–329, doi:10.1002/pro.3786 (2020) PMC6933861.

34 Waterhouse, A. M., Procter, J. B., Martin, D. M., Clamp, M. & Barton, G. J. Jalview Version 2--a multiple sequence alignment editor and analysis workbench. Bioinformatics 25, 1189–1191, doi:10.1093/bioinformatics/btp033 (2009) PMC2672624.

