## Supplementary material for "ADPKD variants in the PKD2 pore helix cause structural collapse of the gate and distinct forms of channel dysfunction": PDB Validation Reports

WORLDWIDE PDB PROTEIN DATA BANK

### Preliminary Full wwPDB EM Validation Report (i)

Aug 27, 2024 – 03:54 PM EDT

Deposition ID : D_1000287792

**This wwPDB validation report is NOT for manuscript review**

This is a Preliminary Full wwPDB EM Validation Report.

This report is produced by the wwPDB Deposition System during initial deposition but before annotation of the structure.

We welcome your comments at **
A user guide is available at
https://www.wwpdb.org/validation/2017/EMValidationReportHelp
with specific help available everywhere you see the (i) symbol.

The types of validation reports are described at
http://www.wwpdb.org/validation/2017/FAQs#types.

Not For Manuscript Review

The following versions of software and data (see references (i)) were used in the production of this report:

| | | |
|---|---|---|
| EMDB validation analysis | : | 0.0.1.dev112 |
| MolProbity | : | 4.02b-467 |
| Percentile statistics | : | 20231227.v01 (using entries in the PDB archive December 27th 2023) |
| MapQ | : | 1.9.13 |
| Ideal geometry (proteins) | : | Engh & Huber (2001) |
| Ideal geometry (DNA, RNA) | : | Parkinson et al. (1996) |
| Validation Pipeline (wwPDB-VP) | : | 2.38.3 |

### 1 Overall quality at a glance (i)

The following experimental techniques were used to determine the structure:
*ELECTRON MICROSCOPY*

The reported resolution of this entry is unknown.

Percentile scores (ranging between 0-100) for global validation metrics of the entry are shown in the following graphic. The table shows the number of entries on which the scores are based.

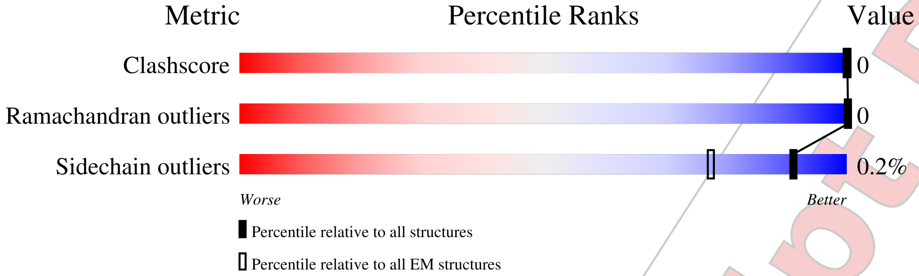


| Metric | Whole archive (#Entries) | EM structures (#Entries) |
|---|---|---|
| Clashscore | 210492 | 15764 |
| Ramachandran outliers | 207382 | 16835 |
| Sidechain outliers | 206894 | 16415 |

The table below summarises the geometric issues observed across the polymeric chains and their fit to the map. The red, orange, yellow and green segments of the bar indicate the fraction of residues that contain outliers for >=3, 2, 1 and 0 types of geometric quality criteria respectively. A grey segment represents the fraction of residues that are not modelled. The numeric value for each fraction is indicated below the corresponding segment, with a dot representing fractions <=5% The upper red bar (where present) indicates the fraction of residues that have poor fit to the EM map (all-atom inclusion < 40%). The numeric value is given above the bar.

| Mol | Chain | Length | Quality of chain |
|---|---|---|---|
| 1 | A1 | 462 | 15% / 98% • |
| 1 | A2 | 462 | 14% / 98% • |
| 1 | A3 | 462 | 14% / 98% • |
| 1 | A4 | 462 | 14% / 98% • |

Not For Manuscript Review

#### 2 Entry composition (i)

There is only 1 type of molecule in this entry. The entry contains 30316 atoms, of which 15080 are hydrogens and 0 are deuteriums.

In the tables below, the AltConf column contains the number of residues with at least one atom in alternate conformation and the Trace column contains the number of residues modelled with at most 2 atoms.

- Molecule 1 is a protein.

| Mol | Chain | Residues | Atoms | | | | | | AltConf | Trace |
|---|---|---|---|---|---|---|---|---|---|---|
| | | | Total | C | H | N | O | S | | |
| 1 | A1 | 462 | 7579 | 2515 | 3770 | 598 | 676 | 20 | 0 | 0 |
| 1 | A2 | 462 | 7579 | 2515 | 3770 | 598 | 676 | 20 | 0 | 0 |
| 1 | A3 | 462 | 7579 | 2515 | 3770 | 598 | 676 | 20 | 0 | 0 |
| 1 | A4 | 462 | 7579 | 2515 | 3770 | 598 | 676 | 20 | 0 | 0 |

Not For Manuscript Review

### 3 Residue-property plots (i)

These plots are drawn for all protein, RNA, DNA and oligosaccharide chains in the entry. The first graphic for a chain summarises the proportions of the various outlier classes displayed in the second graphic. The second graphic shows the sequence view annotated by issues in geometry and atom inclusion in map density. Residues are color-coded according to the number of geometric quality criteria for which they contain at least one outlier: green = 0, yellow = 1, orange = 2 and red = 3 or more. A red diamond above a residue indicates a poor fit to the EM map for this residue (all-atom inclusion < 40%). Stretches of 2 or more consecutive residues without any outlier are shown as a green connector. Residues present in the sample, but not in the model, are shown in grey.

- Molecule 1:

Chain A1:

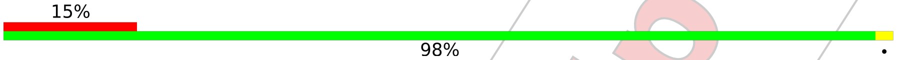


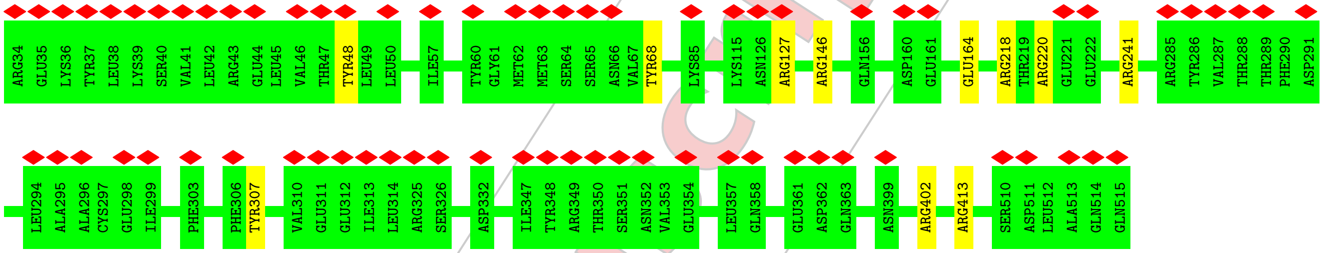


- Molecule 1:

Chain A2:

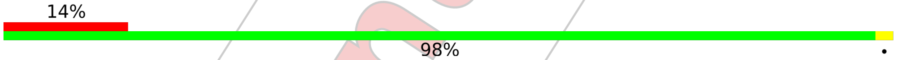


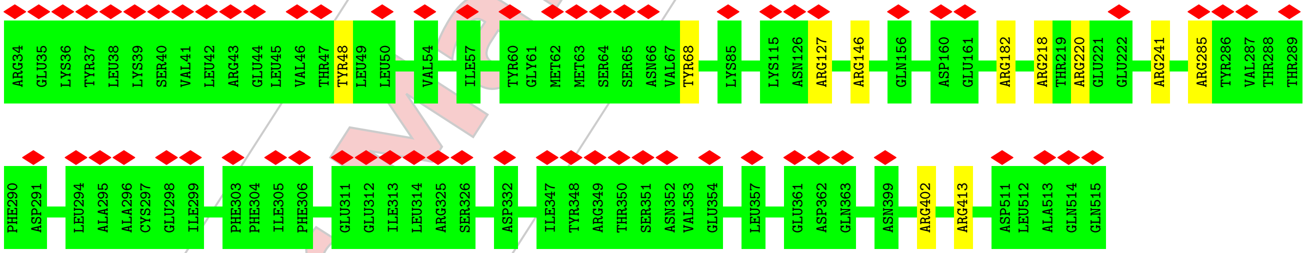


- Molecule 1:

Chain A3:

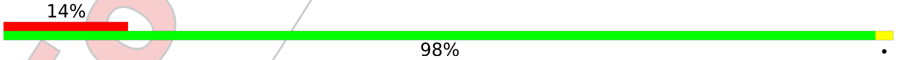


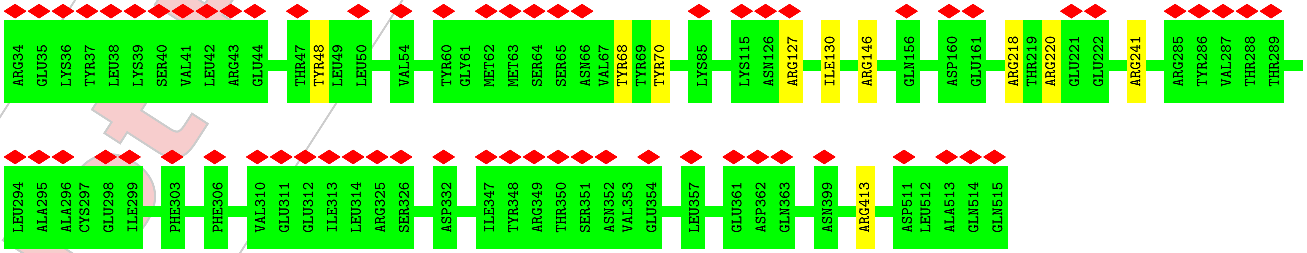


- Molecule 1:

Not For Manuscript Review

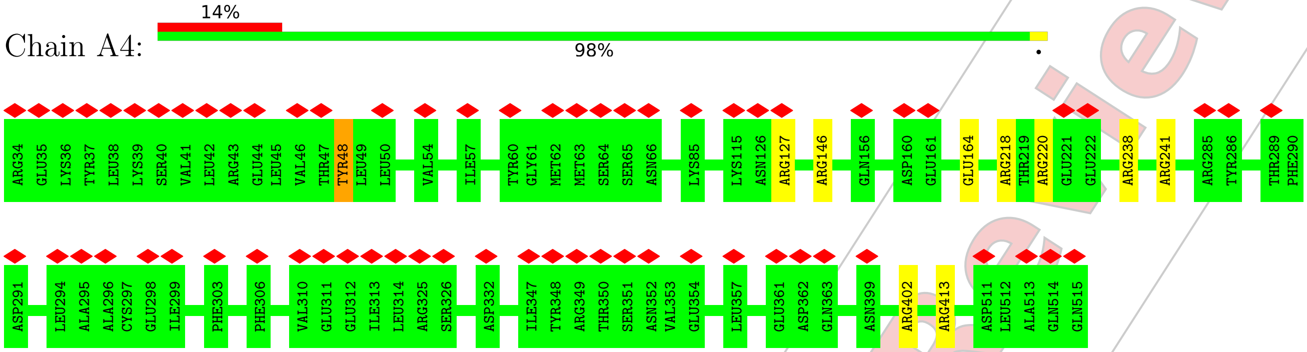


Not For Manuscript Review

### 4 Experimental information (i)

| Property | Value | Source |
|---|---|---|
| EM reconstruction method | SINGLE PARTICLE | Depositor |
| Imposed symmetry | POINT, Not provided | |
| Number of particles used | Not provided | |
| Resolution determination method | Not provided | |
| CTF correction method | Not provided | |
| Microscope | Not provided | |
| Voltage (kV) | Not provided | |
| Electron dose $(e^-/\text{Å}^2)$ | Not provided | |
| Minimum defocus (nm) | Not provided | |
| Maximum defocus (nm) | Not provided | |
| Magnification | Not provided | |
| Image detector | Not provided | |
| Maximum map value | 0.243 | Depositor |
| Minimum map value | -0.198 | Depositor |
| Average map value | 0.000 | Depositor |
| Map value standard deviation | 0.006 | Depositor |
| Recommended contour level | 0.029 | Depositor |
| Map size (Å) | 279.504, 279.504, 279.504 | wwPDB |
| Map dimensions | 864, 864, 864 | wwPDB |
| Map angles (°) | 90.0, 90.0, 90.0 | wwPDB |
| Pixel spacing (Å) | 0.3235, 0.3235, 0.3235 | Depositor |

Not For Manuscript Review

### 5 Model quality (i)

#### 5.1 Standard geometry (i)

The Z score for a bond length (or angle) is the number of standard deviations the observed value is removed from the expected value. A bond length (or angle) with $|Z| > 5$ is considered an outlier worth inspection. RMSZ is the root-mean-square of all Z scores of the bond lengths (or angles).

| Mol | Chain | Bond lengths | | Bond angles | |
|---|---|---|---|---|---|
| | | RMSZ | #$\|Z\|$ >5 | RMSZ | #$\|Z\|$ >5 |
| 1 | A1 | 0.69 | 0/3908 | 0.96 | 8/5302 (0.2%) |
| 1 | A2 | 0.70 | 0/3908 | 0.97 | 10/5302 (0.2%) |
| 1 | A3 | 0.69 | 0/3908 | 0.96 | 8/5302 (0.2%) |
| 1 | A4 | 0.69 | 0/3908 | 0.97 | 9/5302 (0.2%) |
| All | All | 0.69 | 0/15632 | 0.96 | 35/21208 (0.2%) |

Chiral center outliers are detected by calculating the chiral volume of a chiral center and verifying if the center is modelled as a planar moiety or with the opposite hand.A planarity outlier is detected by checking planarity of atoms in a peptide group, atoms in a mainchain group or atoms of a sidechain that are expected to be planar.

| Mol | Chain | #Chirality outliers | #Planarity outliers |
|---|---|---|---|
| 1 | A1 | 0 | 2 |
| 1 | A2 | 0 | 1 |
| 1 | A3 | 0 | 1 |
| 1 | A4 | 0 | 1 |
| All | All | 0 | 5 |

There are no bond length outliers.

All (35) bond angle outliers are listed below:

| Mol | Chain | Res | Type | Atoms | Z | Observed(°) | Ideal(°) |
|---|---|---|---|---|---|---|---|
| 1 | A4 | 241 | ARG | NE-CZ-NH2 | 7.08 | 123.84 | 120.30 |
| 1 | A3 | 220 | ARG | NE-CZ-NH2 | 6.78 | 123.69 | 120.30 |
| 1 | A3 | 241 | ARG | NE-CZ-NH2 | 6.77 | 123.69 | 120.30 |
| 1 | A4 | 413 | ARG | NE-CZ-NH2 | 6.73 | 123.67 | 120.30 |
| 1 | A2 | 413 | ARG | NE-CZ-NH2 | 6.71 | 123.66 | 120.30 |
| 1 | A2 | 241 | ARG | NE-CZ-NH2 | 6.71 | 123.65 | 120.30 |
| 1 | A4 | 48 | TYR | CB-CG-CD1 | -6.71 | 116.97 | 121.00 |
| 1 | A1 | 241 | ARG | NE-CZ-NH2 | 6.71 | 123.65 | 120.30 |
| 1 | A1 | 413 | ARG | NE-CZ-NH2 | 6.70 | 123.65 | 120.30 |
| 1 | A2 | 220 | ARG | NE-CZ-NH2 | 6.67 | 123.63 | 120.30 |

*Continued on next page...*

Not For Manuscript Review

*Continued from previous page...*

| Mol | Chain | Res | Type | Atoms | Z | Observed($^o$) | Ideal($^o$) |
|---|---|---|---|---|---|---|---|
| 1 | A2 | 146 | ARG | NE-CZ-NH2 | 6.62 | 123.61 | 120.30 |
| 1 | A3 | 146 | ARG | NE-CZ-NH2 | 6.58 | 123.59 | 120.30 |
| 1 | A3 | 413 | ARG | NE-CZ-NH2 | 6.51 | 123.56 | 120.30 |
| 1 | A4 | 146 | ARG | NE-CZ-NH2 | 6.43 | 123.51 | 120.30 |
| 1 | A4 | 220 | ARG | NE-CZ-NH2 | 6.40 | 123.50 | 120.30 |
| 1 | A3 | 48 | TYR | CB-CG-CD1 | -6.38 | 117.17 | 121.00 |
| 1 | A1 | 146 | ARG | NE-CZ-NH2 | 6.30 | 123.45 | 120.30 |
| 1 | A2 | 285 | ARG | NE-CZ-NH2 | 6.23 | 123.41 | 120.30 |
| 1 | A2 | 48 | TYR | CB-CG-CD1 | -6.17 | 117.30 | 121.00 |
| 1 | A1 | 220 | ARG | NE-CZ-NH2 | 6.14 | 123.37 | 120.30 |
| 1 | A1 | 48 | TYR | CB-CG-CD1 | -6.09 | 117.35 | 121.00 |
| 1 | A2 | 402 | ARG | NE-CZ-NH2 | 5.95 | 123.28 | 120.30 |
| 1 | A1 | 402 | ARG | NE-CZ-NH2 | 5.89 | 123.24 | 120.30 |
| 1 | A1 | 127 | ARG | NE-CZ-NH2 | 5.76 | 123.18 | 120.30 |
| 1 | A2 | 218 | ARG | NE-CZ-NH2 | 5.72 | 123.16 | 120.30 |
| 1 | A4 | 218 | ARG | NE-CZ-NH2 | 5.70 | 123.15 | 120.30 |
| 1 | A2 | 182 | ARG | NE-CZ-NH2 | 5.67 | 123.14 | 120.30 |
| 1 | A4 | 238 | ARG | NE-CZ-NH2 | 5.60 | 123.10 | 120.30 |
| 1 | A1 | 218 | ARG | NE-CZ-NH2 | 5.55 | 123.07 | 120.30 |
| 1 | A4 | 402 | ARG | NE-CZ-NH2 | 5.49 | 123.04 | 120.30 |
| 1 | A4 | 127 | ARG | NE-CZ-NH2 | 5.34 | 122.97 | 120.30 |
| 1 | A3 | 70 | TYR | CB-CG-CD2 | -5.24 | 117.86 | 121.00 |
| 1 | A2 | 127 | ARG | NE-CZ-NH2 | 5.17 | 122.89 | 120.30 |
| 1 | A3 | 127 | ARG | NE-CZ-NH2 | 5.07 | 122.83 | 120.30 |
| 1 | A3 | 218 | ARG | NE-CZ-NH2 | 5.05 | 122.83 | 120.30 |

There are no chirality outliers.

All (5) planarity outliers are listed below:

| Mol | Chain | Res | Type | Group |
|---|---|---|---|---|
| 1 | A1 | 307 | TYR | Sidechain |
| 1 | A1 | 68 | TYR | Sidechain |
| 1 | A2 | 68 | TYR | Sidechain |
| 1 | A3 | 68 | TYR | Sidechain |
| 1 | A4 | 48 | TYR | Sidechain |

#### 5.2 Too-close contacts ⓘ

In the following table, the Non-H and H(model) columns list the number of non-hydrogen atoms and hydrogen atoms in the chain respectively. The H(added) column lists the number of hydrogen atoms added and optimized by MolProbity. The Clashes column lists the number of clashes within

Not For Manuscript Review

the asymmetric unit, whereas Symm-Clashes lists symmetry-related clashes.

| Mol | Chain | Non-H | H(model) | H(added) | Clashes | Symm-Clashes |
|---|---|---|---|---|---|---|
| 1 | A1 | 3809 | 3770 | 3763 | 0 | 0 |
| 1 | A2 | 3809 | 3770 | 3763 | 0 | 0 |
| 1 | A3 | 3809 | 3770 | 3763 | 0 | 0 |
| 1 | A4 | 3809 | 3770 | 3763 | 0 | 0 |
| All | All | 15236 | 15080 | 15052 | 0 | 0 |

The all-atom clashscore is defined as the number of clashes found per 1000 atoms (including hydrogen atoms). The all-atom clashscore for this structure is 0.

There are no clashes within the asymmetric unit.

There are no symmetry-related clashes.

#### 5.3 Torsion angles (i)

##### 5.3.1 Protein backbone (i)

In the following table, the Percentiles column shows the percent Ramachandran outliers of the chain as a percentile score with respect to all PDB entries followed by that with respect to all EM entries.

The Analysed column shows the number of residues for which the backbone conformation was analysed, and the total number of residues.

| Mol | Chain | Analysed | Favoured | Allowed | Outliers | Percentiles | |
|---|---|---|---|---|---|---|---|
| 1 | A1 | 456/462 (99%) | 439 (96%) | 17 (4%) | 0 | 100 | 100 |
| 1 | A2 | 456/462 (99%) | 434 (95%) | 22 (5%) | 0 | 100 | 100 |
| 1 | A3 | 456/462 (99%) | 437 (96%) | 19 (4%) | 0 | 100 | 100 |
| 1 | A4 | 456/462 (99%) | 438 (96%) | 18 (4%) | 0 | 100 | 100 |
| All | All | 1824/1848 (99%) | 1748 (96%) | 76 (4%) | 0 | 100 | 100 |

There are no Ramachandran outliers to report.

##### 5.3.2 Protein sidechains (i)

In the following table, the Percentiles column shows the percent sidechain outliers of the chain as a percentile score with respect to all PDB entries followed by that with respect to all EM entries.

The Analysed column shows the number of residues for which the sidechain conformation was analysed, and the total number of residues.

Not For Manuscript Review

| Mol | Chain | Analysed | Rotameric | Outliers | Percentiles | |
|---|---|---|---|---|---|---|
| 1 | A1 | 418/418 (100%) | 417 (100%) | 1 (0%) | 92 | 92 |
| 1 | A2 | 418/418 (100%) | 418 (100%) | 0 | 100 | 100 |
| 1 | A3 | 418/418 (100%) | 417 (100%) | 1 (0%) | 92 | 92 |
| 1 | A4 | 418/418 (100%) | 417 (100%) | 1 (0%) | 92 | 92 |
| All | All | 1672/1672 (100%) | 1669 (100%) | 3 (0%) | 91 | 92 |

All (3) residues with a non-rotameric sidechain are listed below:

| Mol | Chain | Res | Type |
|---|---|---|---|
| 1 | A1 | 164 | GLU |
| 1 | A3 | 130 | ILE |
| 1 | A4 | 164 | GLU |

Sometimes sidechains can be flipped to improve hydrogen bonding and reduce clashes. There are no such sidechains identified.

##### 5.3.3 RNA (i)

There are no RNA molecules in this entry.

#### 5.4 Non-standard residues in protein, DNA, RNA chains (i)

There are no non-standard protein/DNA/RNA residues in this entry.

#### 5.5 Carbohydrates (i)

There are no oligosaccharides in this entry.

#### 5.6 Ligand geometry (i)

There are no ligands in this entry.

#### 5.7 Other polymers (i)

There are no such residues in this entry.

Not For Manuscript Review

#### 5.8 Polymer linkage issues (i)

The following chains have linkage breaks:

| Mol | Chain | Number of breaks |
|---|---|---|
| 1 | A3 | 2 |
| 1 | A4 | 2 |
| 1 | A1 | 2 |
| 1 | A2 | 2 |

All chain breaks are listed below:

| Model | Chain | Residue-1 | Atom-1 | Residue-2 | Atom-2 | Distance (Å) |
|---|---|---|---|---|---|---|
| 1 | A3 | 314:LEU | C | 325:ARG | N | 14.28 |
| 1 | A4 | 314:LEU | C | 325:ARG | N | 13.97 |
| 1 | A1 | 314:LEU | C | 325:ARG | N | 13.84 |
| 1 | A2 | 314:LEU | C | 325:ARG | N | 13.19 |
| 1 | A1 | 115:LYS | C | 126:ASN | N | 10.79 |
| 1 | A4 | 115:LYS | C | 126:ASN | N | 10.77 |
| 1 | A3 | 115:LYS | C | 126:ASN | N | 10.75 |
| 1 | A2 | 115:LYS | C | 126:ASN | N | 10.33 |

Not For Manuscript Review

### 6 Map visualisation (i)

This section contains visualisations of the EMDB entry D_1000287792. These allow visual inspection of the internal detail of the map and identification of artifacts.

Images derived from a raw map, generated by summing the deposited half-maps, are presented below the corresponding image components of the primary map to allow further visual inspection and comparison with those of the primary map.

#### 6.1 Orthogonal projections (i)

##### 6.1.1 Primary map

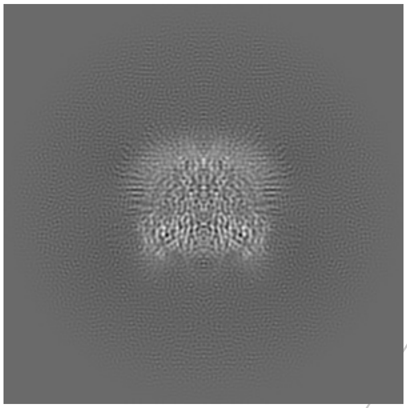

X

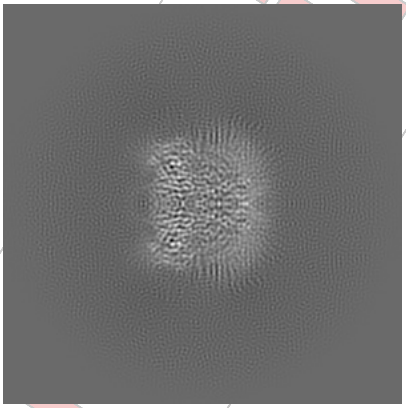

Y

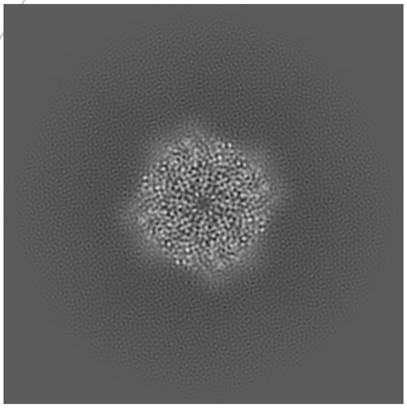

Z

##### 6.1.2 Raw map

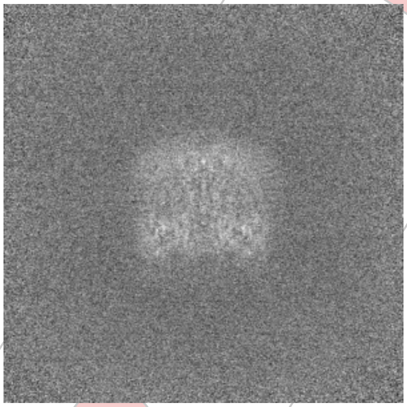

X

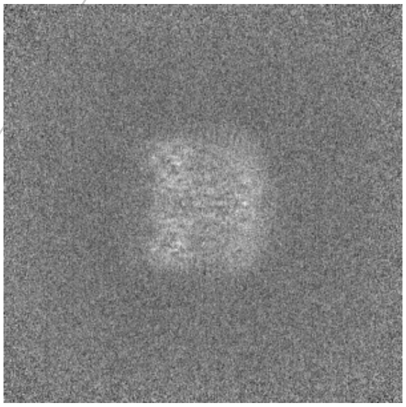

Y

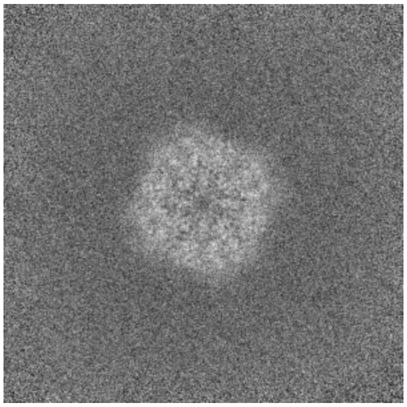

Z

The images above show the map projected in three orthogonal directions.

Not For Manuscript Review

#### 6.2 Central slices (i)

##### 6.2.1 Primary map

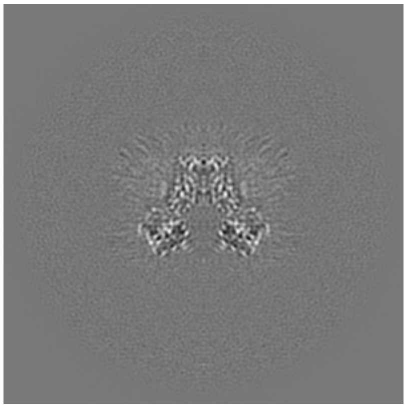
X Index: 432

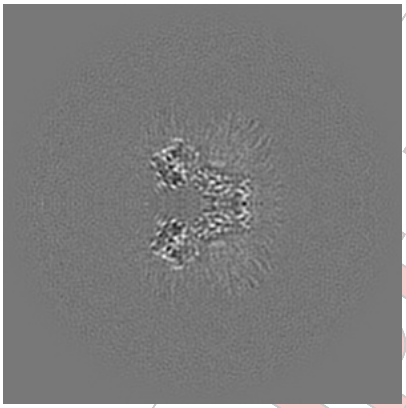
Y Index: 432

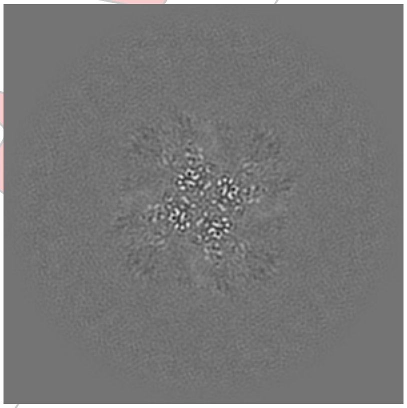
Z Index: 432

##### 6.2.2 Raw map

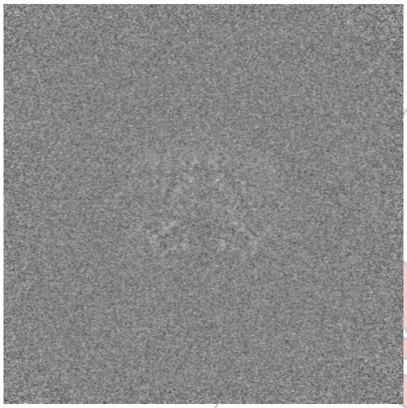
X Index: 432

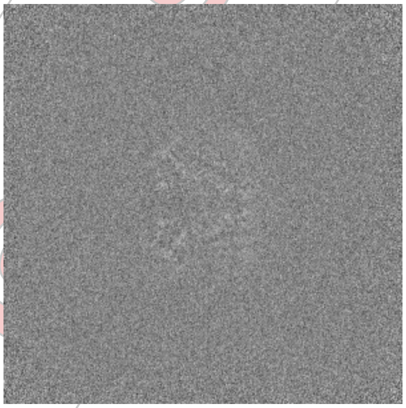
Y Index: 432

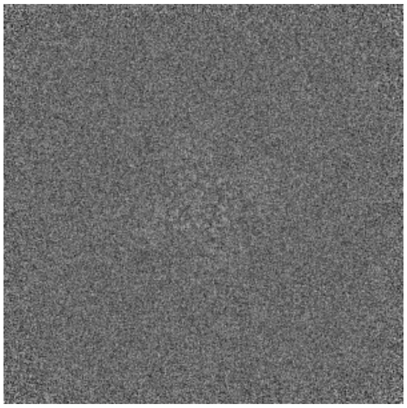
Z Index: 432

The images above show central slices of the map in three orthogonal directions.

Not For Manuscript Review

#### 6.3 Largest variance slices (i)

##### 6.3.1 Primary map

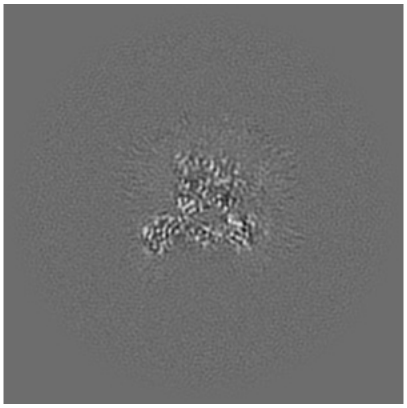

X Index: 387

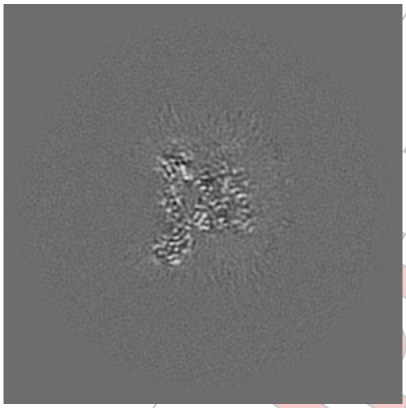

Y Index: 477

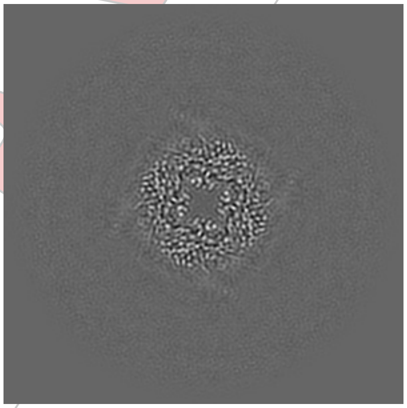

Z Index: 372

##### 6.3.2 Raw map

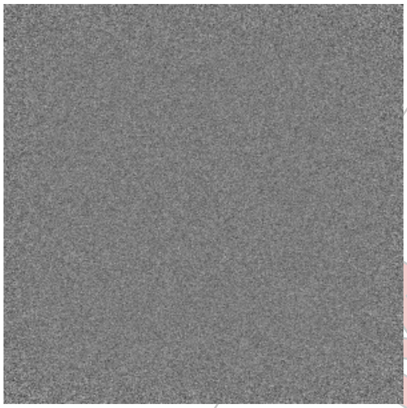

X Index: 0

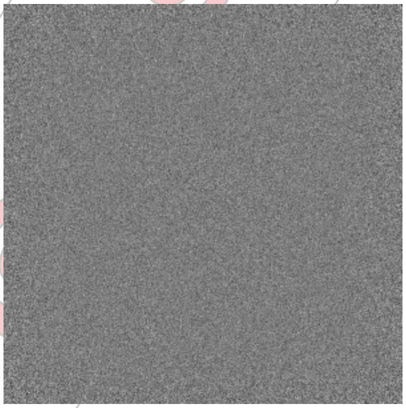

Y Index: 0

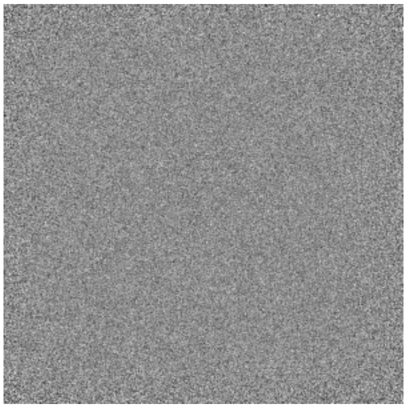

Z Index: 0

The images above show the largest variance slices of the map in three orthogonal directions.

Not For Manuscript Review

#### 6.4 Orthogonal standard-deviation projections (False-color) (i)

##### 6.4.1 Primary map

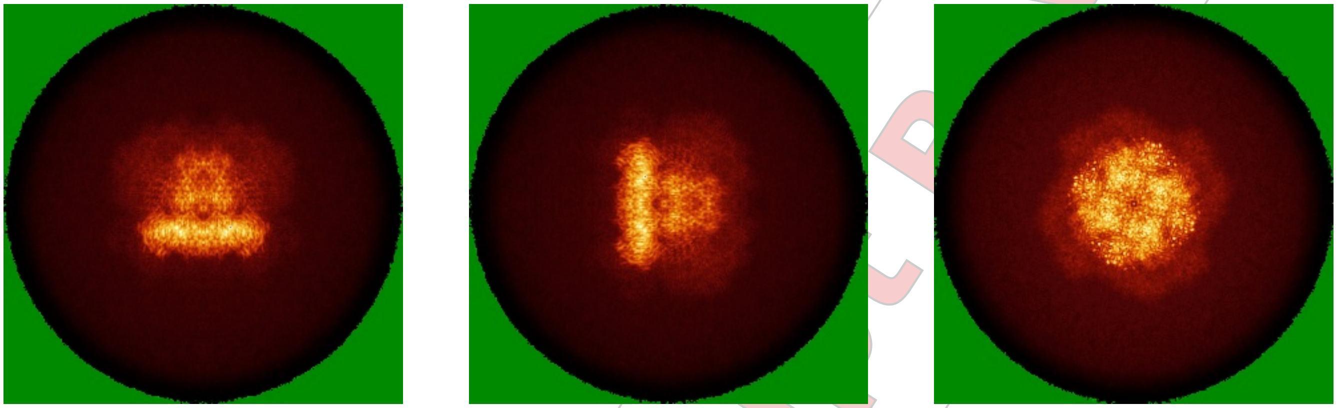

X Y Z

##### 6.4.2 Raw map

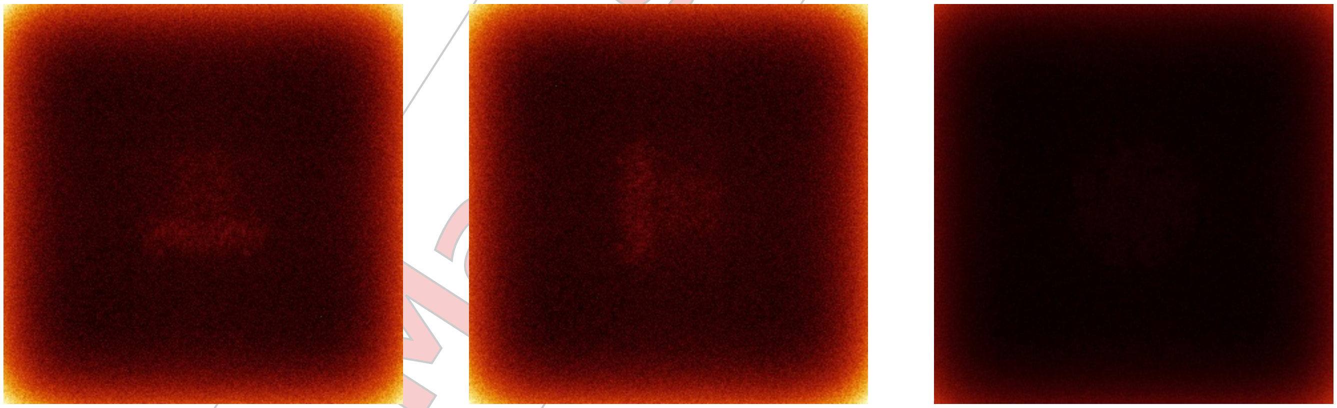

X Y Z

The images above show the map standard deviation projections with false color in three orthogonal directions. Minimum values are shown in green, max in blue, and dark to light orange shades represent small to large values respectively.

Not For Manuscript Review

#### 6.5 Orthogonal surface views (i)

##### 6.5.1 Primary map

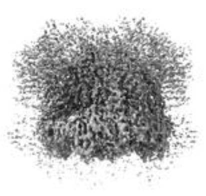

X

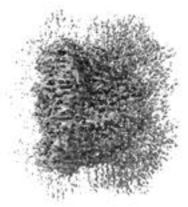

Y

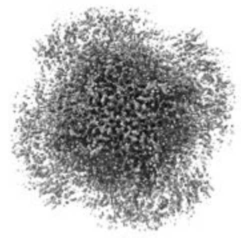

Z

The images above show the 3D surface view of the map at the recommended contour level 0.029. These images, in conjunction with the slice images, may facilitate assessment of whether an appropriate contour level has been provided.

##### 6.5.2 Raw map

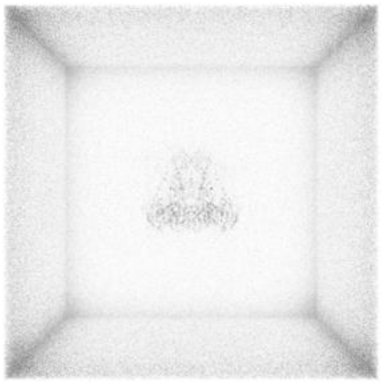

X

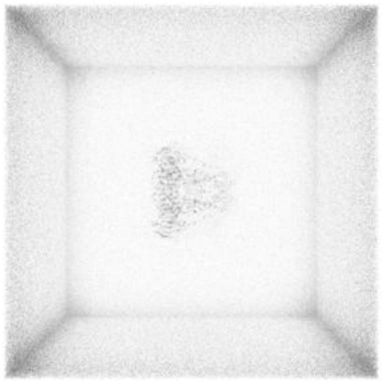

Y

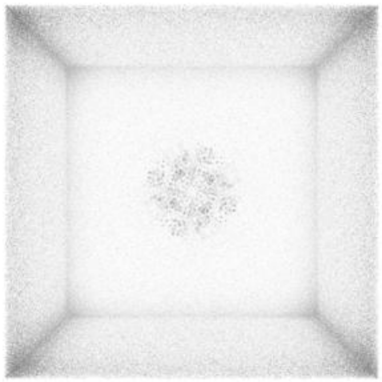

Z

These images show the 3D surface of the raw map. The raw map's contour level was selected so that its surface encloses the same volume as the primary map does at its recommended contour level.

Not For Manuscript Review

#### 6.6 Mask visualisation (i)

This section shows the 3D surface view of the primary map at 50% transparency overlaid with the specified mask at 0% transparency

A mask typically either:

- Encompasses the whole structure
- Separates out a domain, a functional unit, a monomer or an area of interest from a larger structure

##### 6.6.1 D_1000287792_em-mask-volume_P1.map.V2 (i)

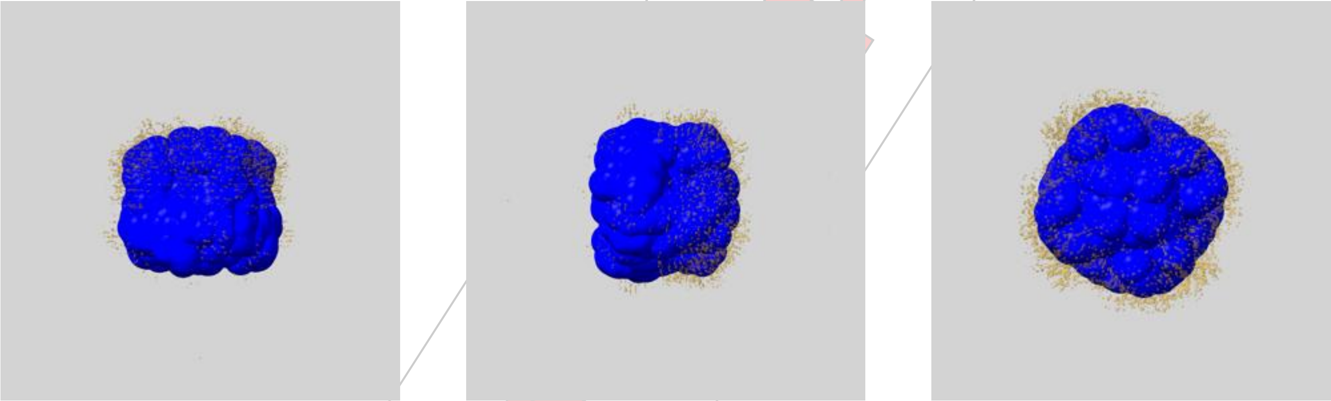

X Y Z

Not For Manuscript Review

### 7 Map analysis (i)

This section contains the results of statistical analysis of the map.

#### 7.1 Map-value distribution (i)

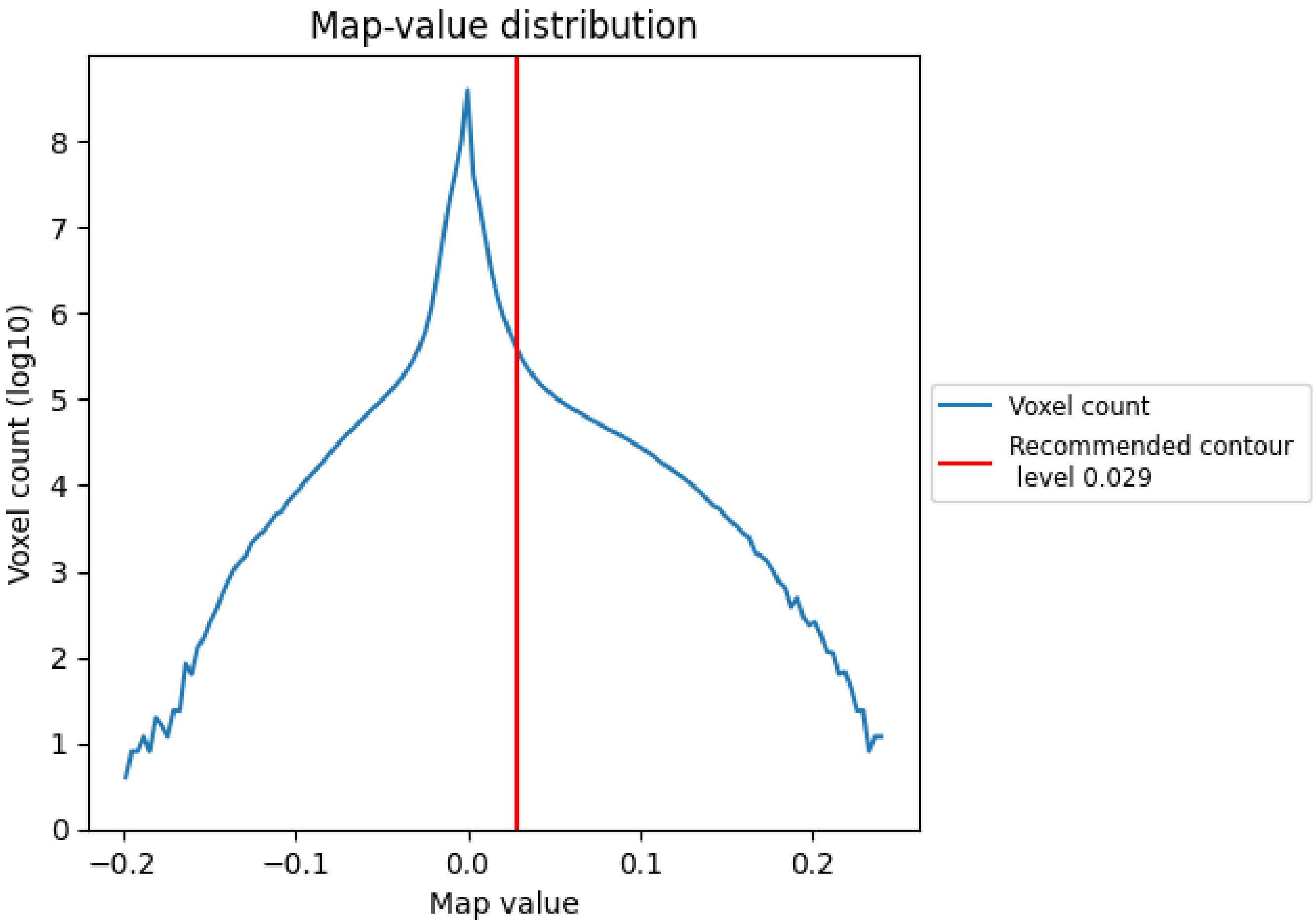


The map-value distribution is plotted in 128 intervals along the x-axis. The y-axis is logarithmic. A spike in this graph at zero usually indicates that the volume has been masked.

Not For Manuscript Review

#### 7.2 Volume estimate (i)

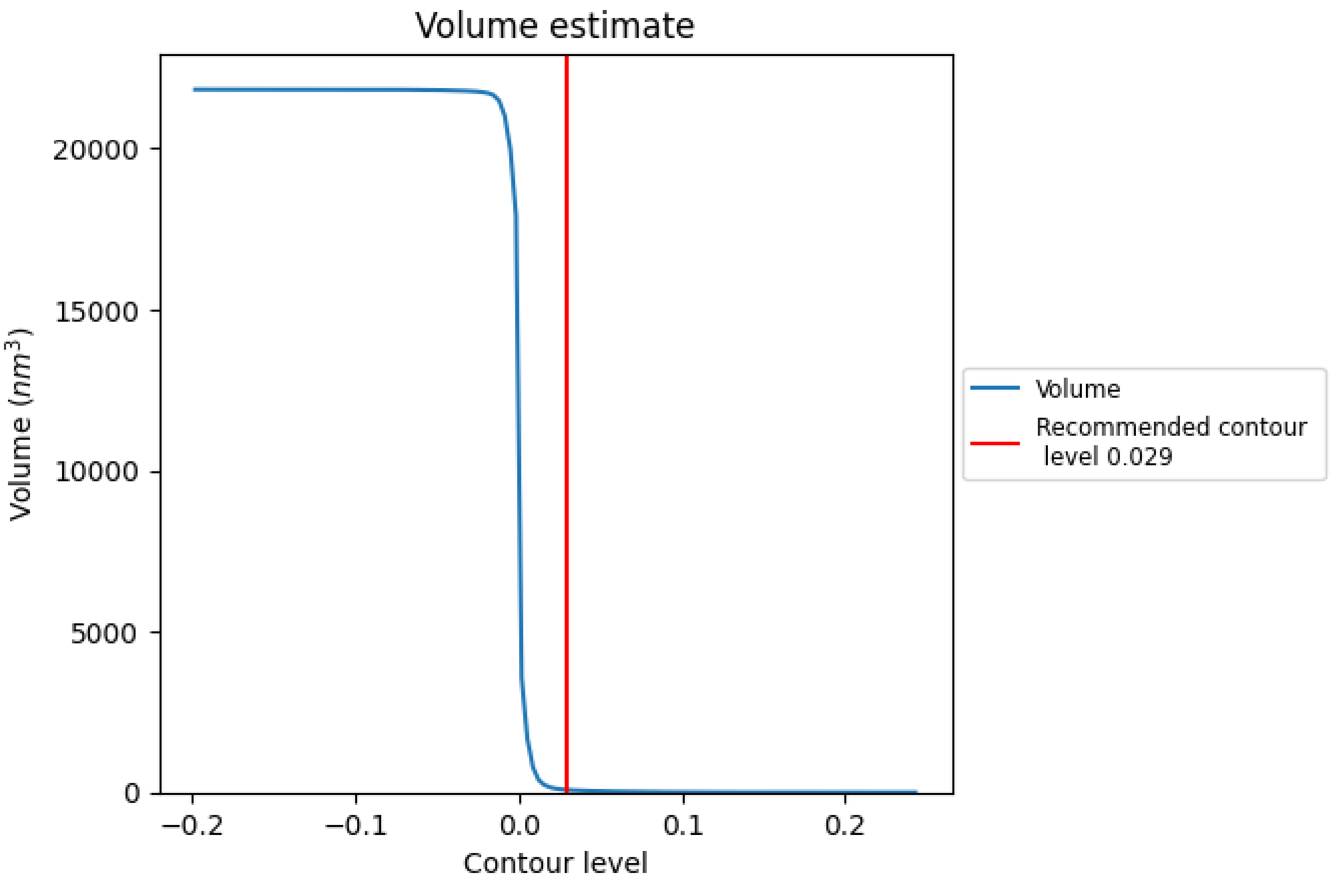


The volume at the recommended contour level is 81 $nm^3$; this corresponds to an approximate mass of 73 kDa.

The volume estimate graph shows how the enclosed volume varies with the contour level. The recommended contour level is shown as a vertical line and the intersection between the line and the curve gives the volume of the enclosed surface at the given level.

Not For Manuscript Review

#### 7.3 Rotationally averaged power spectrum (i)

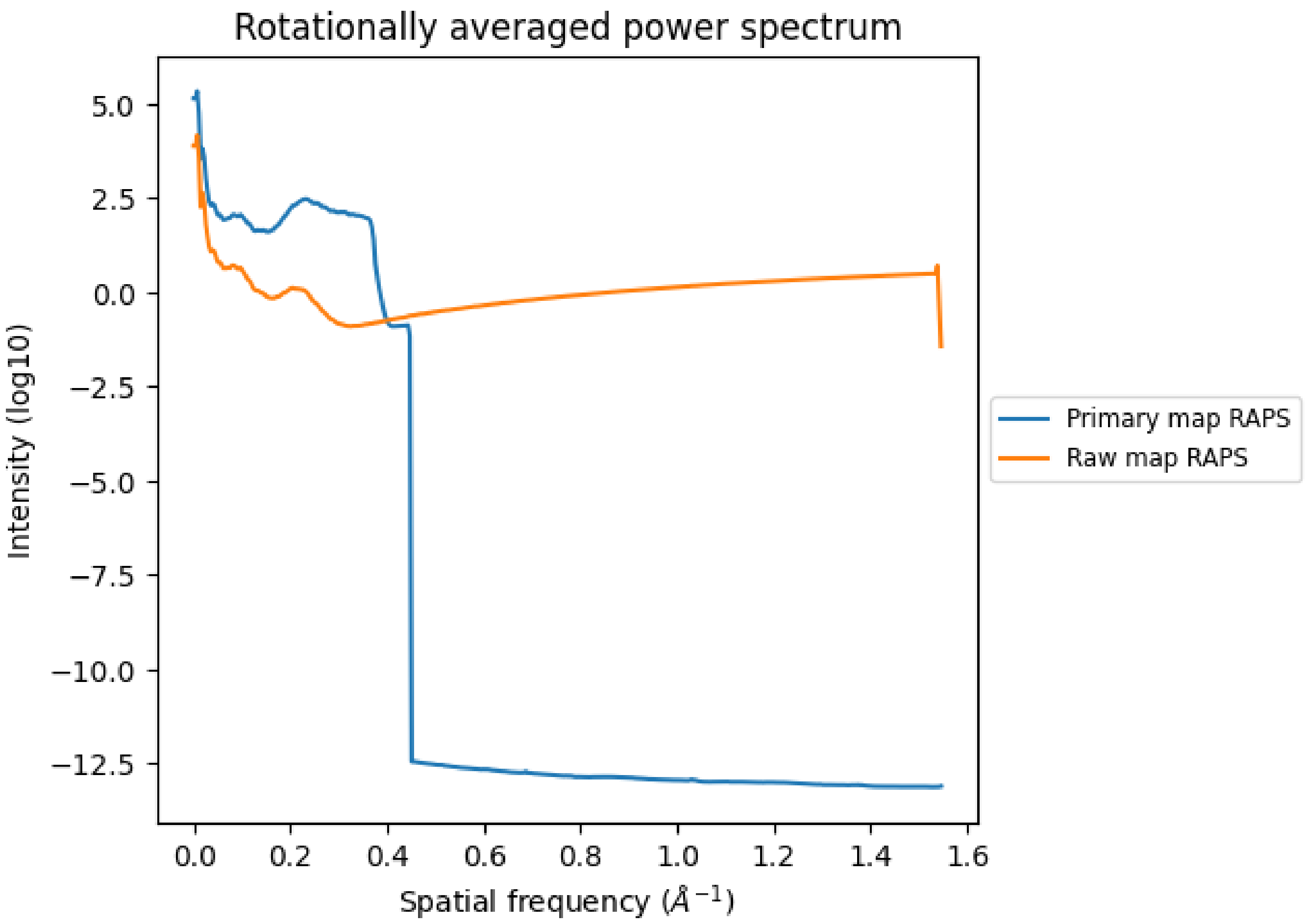


Not For Manuscript Review

### 8 Fourier-Shell correlation (i)

Fourier-Shell Correlation (FSC) is the most commonly used method to estimate the resolution of single-particle and subtomogram-averaged maps. The shape of the curve depends on the imposed symmetry, mask and whether or not the two 3D reconstructions used were processed from a common reference. The reported resolution is shown as a black line. A curve is displayed for the half-bit criterion in addition to lines showing the 0.143 gold standard cut-off and 0.5 cut-off.

#### 8.1 FSC (i)

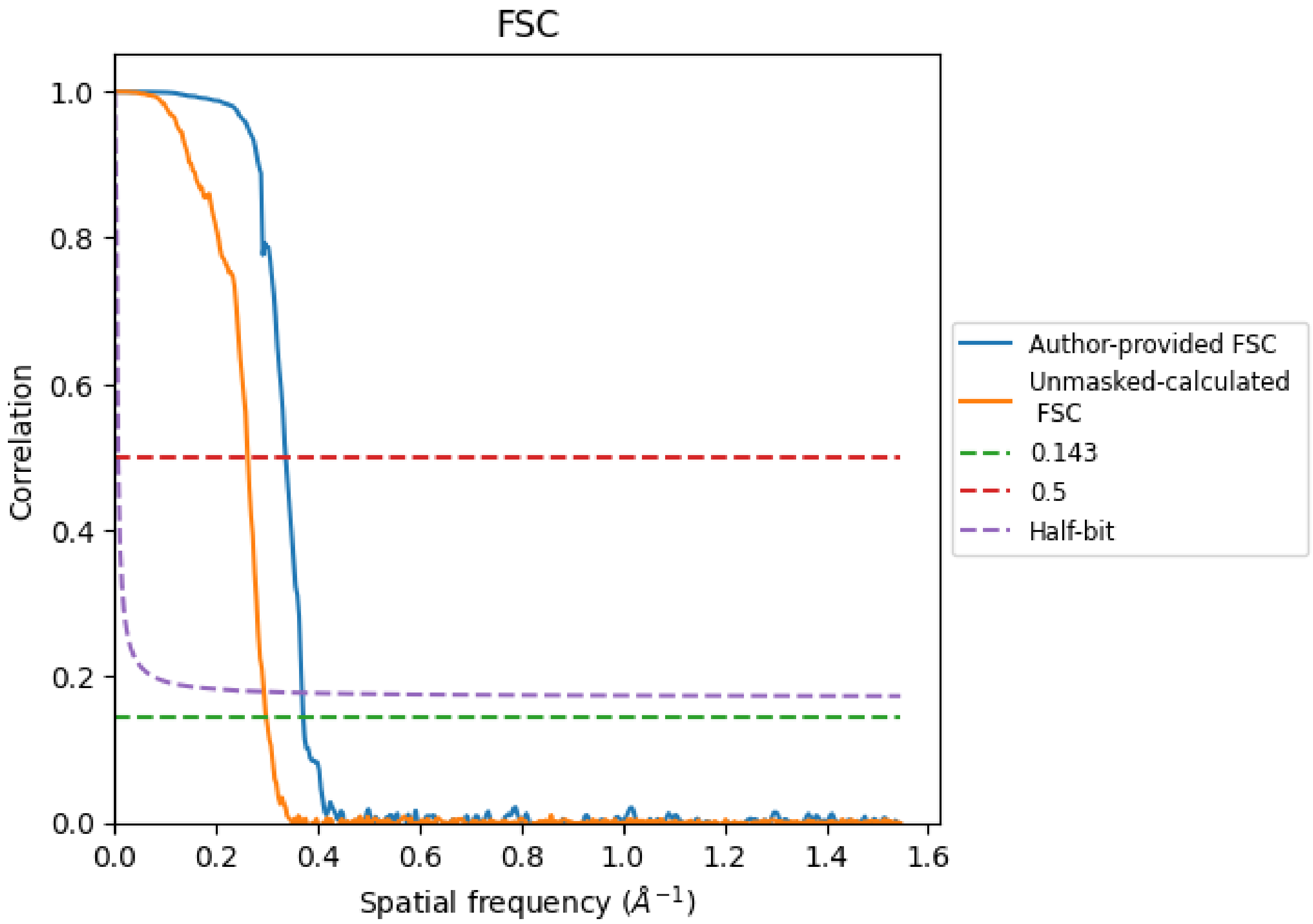


Not For Manuscript Review

#### 8.2 Resolution estimates (i)

| Resolution estimate (Å) | Estimation criterion (FSC cut-off) | | |
|---|---|---|---|
| | 0.143 | 0.5 | Half-bit |
| Reported by author | - | - | - |
| Author-provided FSC curve | 2.69 | 2.97 | 2.71 |
| Unmasked-calculated* | 3.35 | 3.82 | 3.41 |

*Resolution estimate based on FSC curve calculated by comparison of deposited half-maps.

Not For Manuscript Review

### 9 Map-model fit (i)

This section contains information regarding the fit between EMDB map D_1000287792 and PDB model D_1000287792. Per-residue inclusion information can be found in section 3 on page 4.

#### 9.1 Map-model overlay (i)

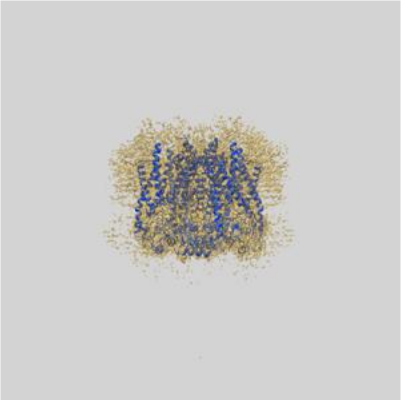

X

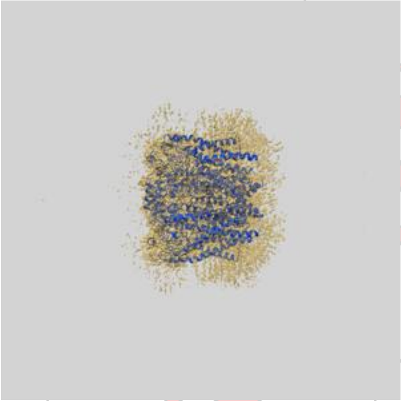

Y

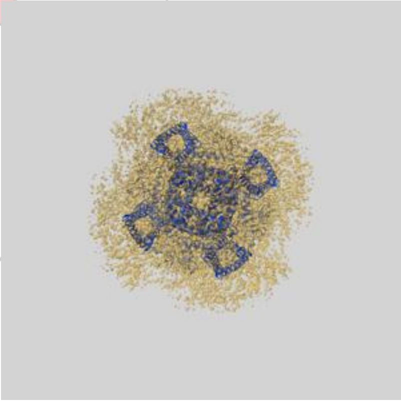

Z

The images above show the 3D surface view of the map at the recommended contour level 0.029 at 50% transparency in yellow overlaid with a ribbon representation of the model coloured in blue. These images allow for the visual assessment of the quality of fit between the atomic model and the map.

Not For Manuscript Review

#### 9.2 Q-score mapped to coordinate model (i)

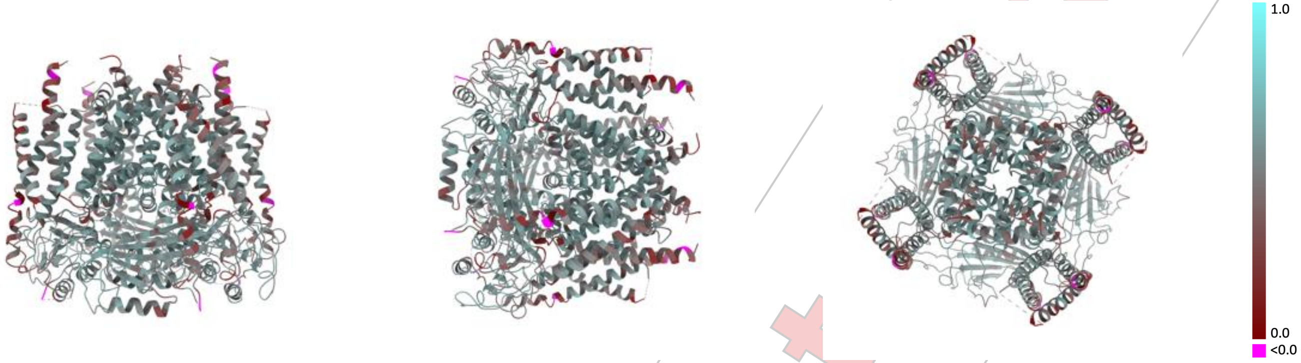


X Y Z

The images above show the model with each residue coloured according its Q-score. This shows their resolvability in the map with higher Q-score values reflecting better resolvability. Please note: Q-score is calculating the resolvability of atoms, and thus high values are only expected at resolutions at which atoms can be resolved. Low Q-score values may therefore be expected for many entries.

#### 9.3 Atom inclusion mapped to coordinate model (i)

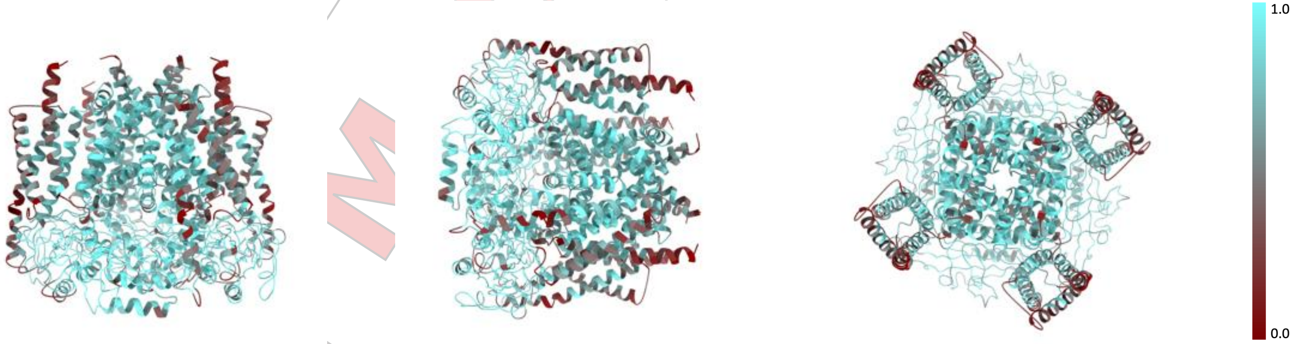


X Y Z

The images above show the model with each residue coloured according to its atom inclusion. This shows to what extent they are inside the map at the recommended contour level (0.029).

Not For Manuscript Review

#### 9.4 Atom inclusion (i)

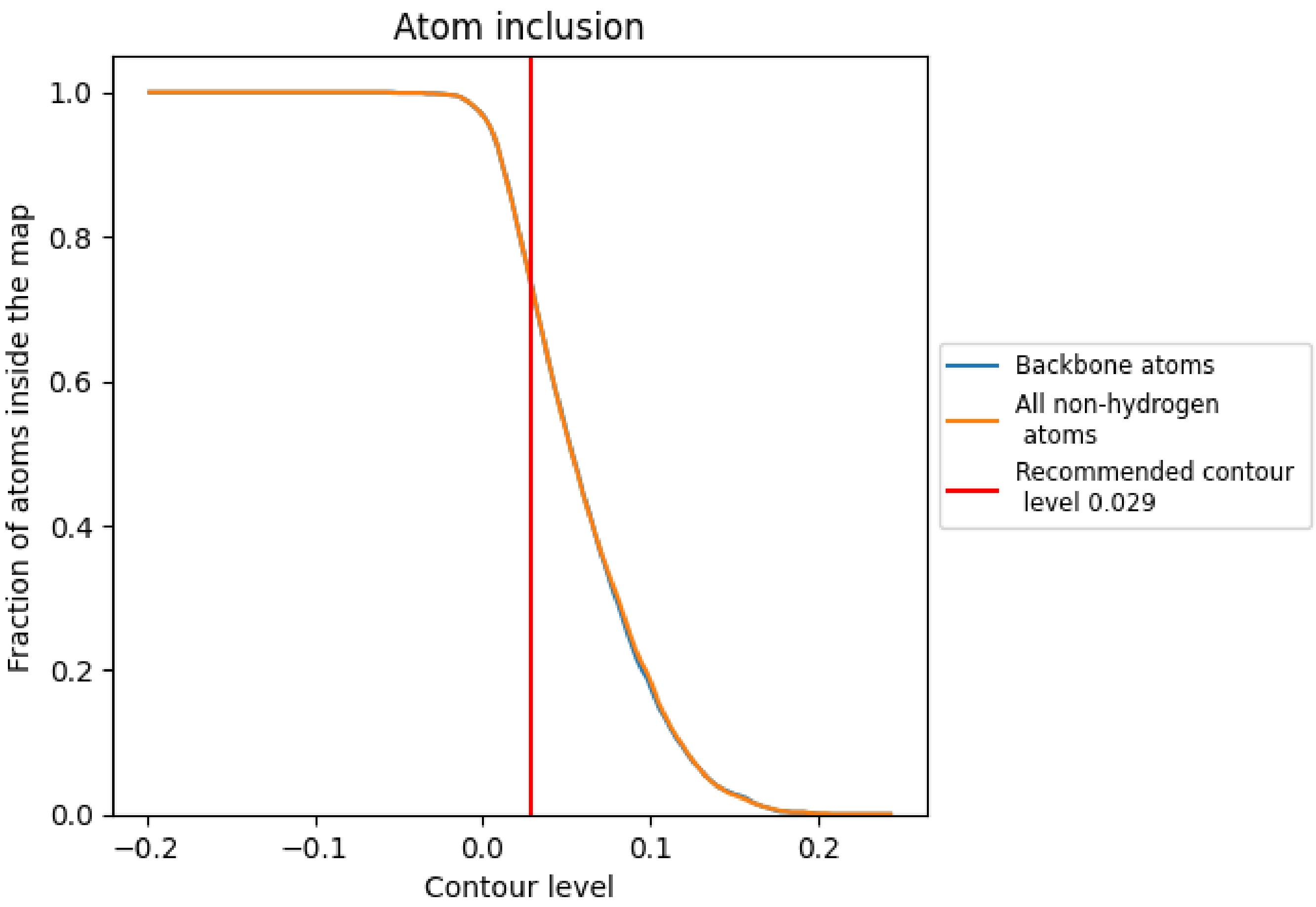


At the recommended contour level, 74% of all backbone atoms, 73% of all non-hydrogen atoms, are inside the map.

Not For Manuscript Review

#### 9.5 Map-model fit summary (i)

The table lists the average atom inclusion at the recommended contour level (0.029) and Q-score for the entire model and for each chain.

| Chain | Atom inclusion | Q-score |
|---|---|---|
| All | 0.7340 | 0.5000 |
| A1 | 0.7360 | 0.5010 |
| A2 | 0.7350 | 0.5000 |
| A3 | 0.7380 | 0.5000 |
| A4 | 0.7360 | 0.5010 |

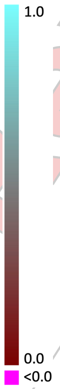


Not For Manuscript Review

### Preliminary Full wwPDB EM Validation Report (i)

Aug 29, 2024 – 02:32 PM EDT

Deposition ID : D_1000287791

**This wwPDB validation report is NOT for manuscript review**

This is a Preliminary Full wwPDB EM Validation Report.

This report is produced by the wwPDB Deposition System
during initial deposition but before annotation of the structure.

We welcome your comments at **
A user guide is available at
https://www.wwpdb.org/validation/2017/EMValidationReportHelp
with specific help available everywhere you see the (i) symbol.

The types of validation reports are described at
http://www.wwpdb.org/validation/2017/FAQs#types.

Not For Manuscript Review

The following versions of software and data (see references (i)) were used in the production of this report:

| | | |
|---|---|---|
| EMDB validation analysis | : | 0.0.1.dev112 |
| MolProbity | : | 4.02b-467 |
| Percentile statistics | : | 20231227.v01 (using entries in the PDB archive December 27th 2023) |
| MapQ | : | 1.9.13 |
| Ideal geometry (proteins) | : | Engh & Huber (2001) |
| Ideal geometry (DNA, RNA) | : | Parkinson et al. (1996) |
| Validation Pipeline (wwPDB-VP) | : | 2.38.3 |

### 1 Overall quality at a glance (i)

The following experimental techniques were used to determine the structure:
*ELECTRON MICROSCOPY*

The reported resolution of this entry is unknown.

Percentile scores (ranging between 0-100) for global validation metrics of the entry are shown in the following graphic. The table shows the number of entries on which the scores are based.

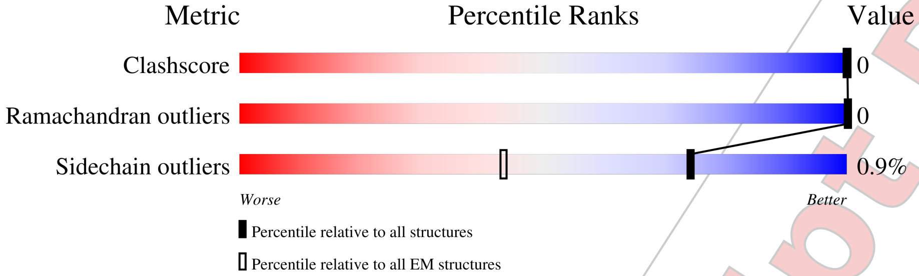


| Metric | Whole archive (#Entries) | EM structures (#Entries) |
|---|---|---|
| Clashscore | 210492 | 15764 |
| Ramachandran outliers | 207382 | 16835 |
| Sidechain outliers | 206894 | 16415 |

The table below summarises the geometric issues observed across the polymeric chains and their fit to the map. The red, orange, yellow and green segments of the bar indicate the fraction of residues that contain outliers for >=3, 2, 1 and 0 types of geometric quality criteria respectively. A grey segment represents the fraction of residues that are not modelled. The numeric value for each fraction is indicated below the corresponding segment, with a dot representing fractions <=5% The upper red bar (where present) indicates the fraction of residues that have poor fit to the EM map (all-atom inclusion < 40%). The numeric value is given above the bar.

| Mol | Chain | Length | Quality of chain |
|---|---|---|---|
| 1 | A | 462 | 26% / 97% • |
| 1 | B | 462 | 25% / 96% • |
| 1 | C | 462 | 25% / 96% • |
| 1 | D | 462 | 26% / 95% 5% |

Not For Manuscript Review

#### 2 Entry composition (i)

There is only 1 type of molecule in this entry. The entry contains 30336 atoms, of which 15088 are hydrogens and 0 are deuteriums.

In the tables below, the AltConf column contains the number of residues with at least one atom in alternate conformation and the Trace column contains the number of residues modelled with at most 2 atoms.

- Molecule 1 is a protein.

| Mol | Chain | Residues | Atoms | | | | | | AltConf | Trace |
|---|---|---|---|---|---|---|---|---|---|---|
| | | | Total | C | H | N | O | S | | |
| 1 | A | 462 | 7584 | 2516 | 3772 | 601 | 676 | 19 | 0 | 0 |
| 1 | B | 462 | 7584 | 2516 | 3772 | 601 | 676 | 19 | 0 | 0 |
| 1 | C | 462 | 7584 | 2516 | 3772 | 601 | 676 | 19 | 0 | 0 |
| 1 | D | 462 | 7584 | 2516 | 3772 | 601 | 676 | 19 | 0 | 0 |

Not For Manuscript Review

### 3 Residue-property plots ⓘ

These plots are drawn for all protein, RNA, DNA and oligosaccharide chains in the entry. The first graphic for a chain summarises the proportions of the various outlier classes displayed in the second graphic. The second graphic shows the sequence view annotated by issues in geometry and atom inclusion in map density. Residues are color-coded according to the number of geometric quality criteria for which they contain at least one outlier: green = 0, yellow = 1, orange = 2 and red = 3 or more. A red diamond above a residue indicates a poor fit to the EM map for this residue (all-atom inclusion < 40%). Stretches of 2 or more consecutive residues without any outlier are shown as a green connector. Residues present in the sample, but not in the model, are shown in grey.

- Molecule 1:

Chain A:

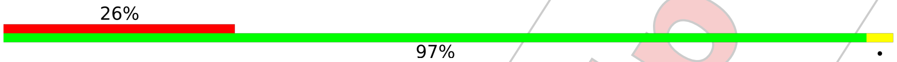


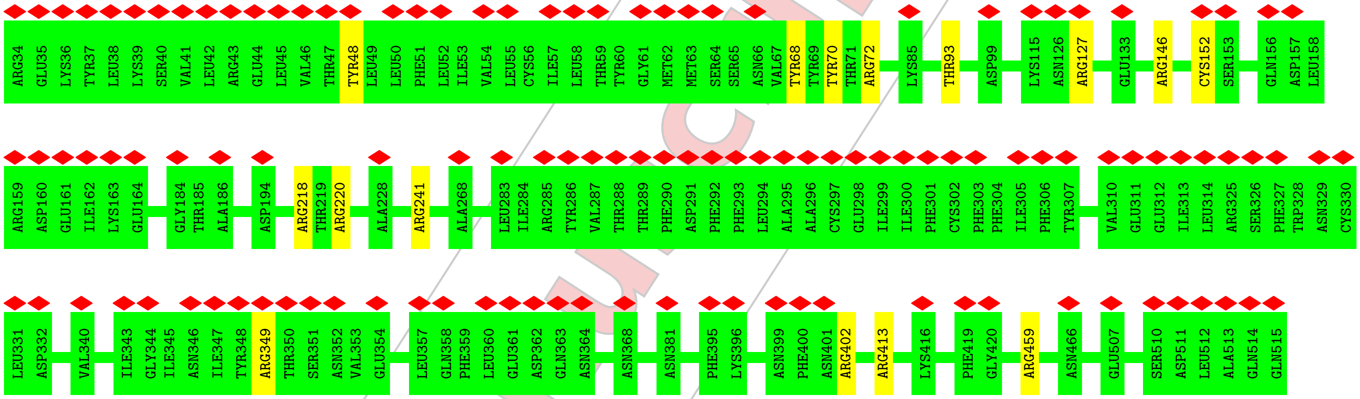


- Molecule 1:

Chain B:

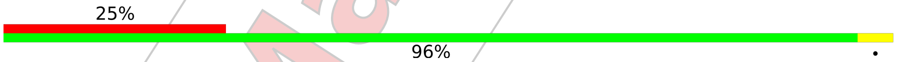


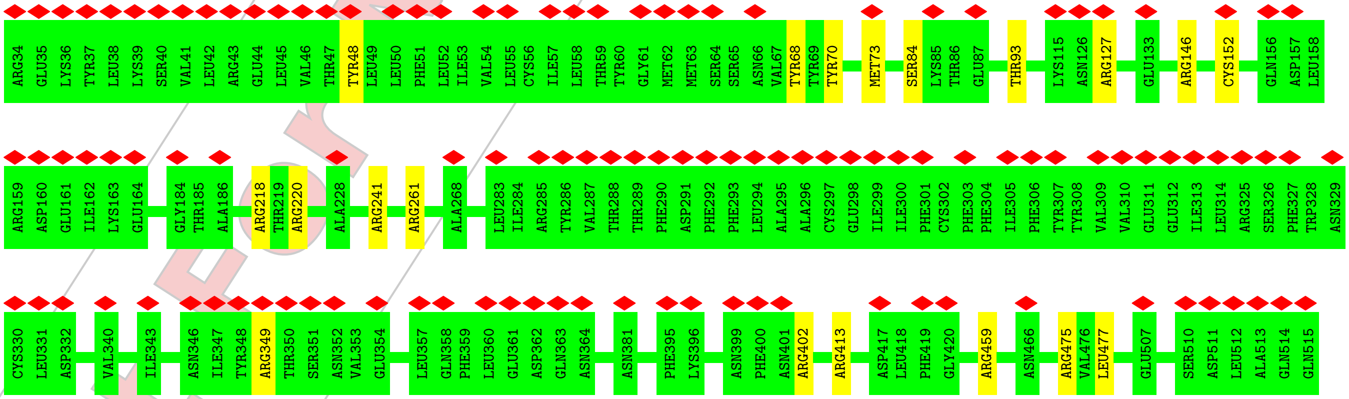


- Molecule 1:

Chain C:

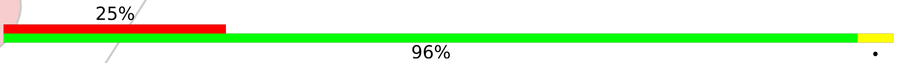


Not For Manuscript Review

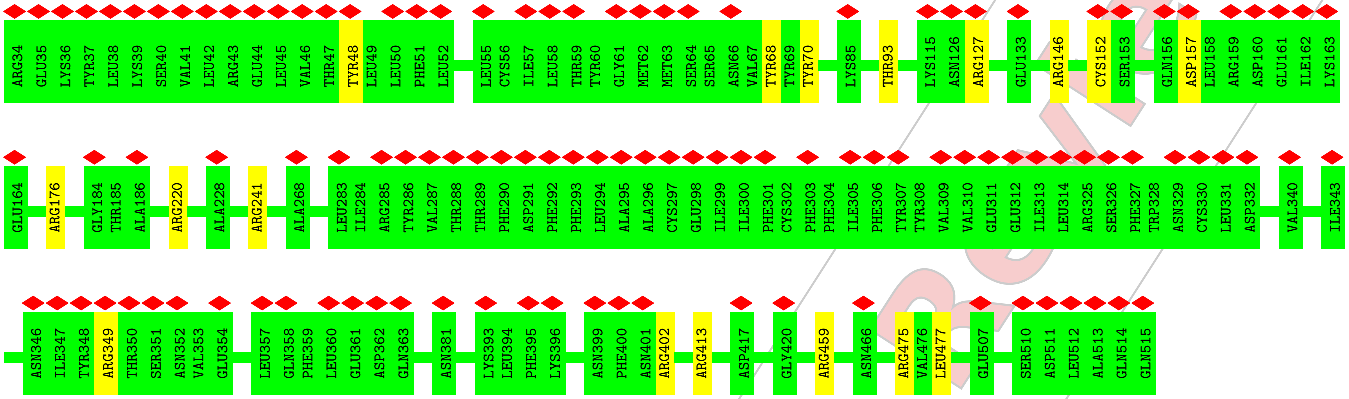


- Molecule 1:

Chain D:

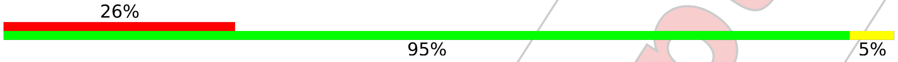


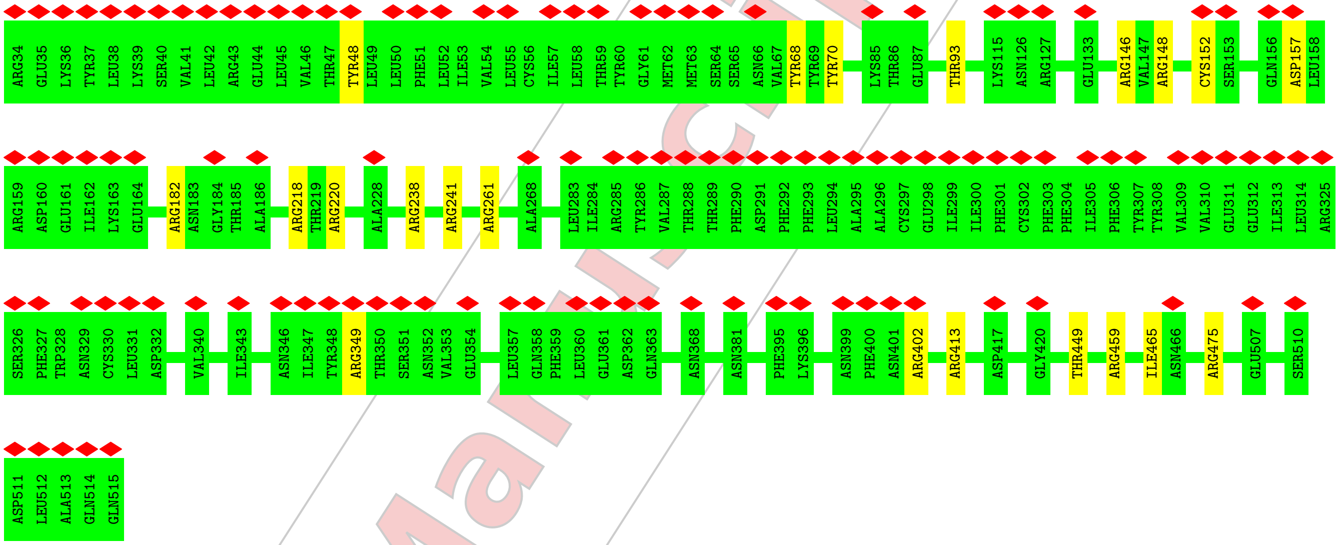


Not For Manuscript Review

### 4 Experimental information (i)

| Property | Value | Source |
|---|---|---|
| EM reconstruction method | SINGLE PARTICLE | Depositor |
| Imposed symmetry | POINT, Not provided | |
| Number of particles used | Not provided | |
| Resolution determination method | Not provided | |
| CTF correction method | Not provided | |
| Microscope | Not provided | |
| Voltage (kV) | Not provided | |
| Electron dose $(e^-/\text{Å}^2)$ | Not provided | |
| Minimum defocus (nm) | Not provided | |
| Maximum defocus (nm) | Not provided | |
| Magnification | Not provided | |
| Image detector | Not provided | |
| Maximum map value | 0.443 | Depositor |
| Minimum map value | -0.296 | Depositor |
| Average map value | 0.000 | Depositor |
| Map value standard deviation | 0.015 | Depositor |
| Recommended contour level | 0.0891 | Depositor |
| Map size (Å) | 258.79825, 258.79825, 258.79825 | wwPDB |
| Map dimensions | 432, 432, 432 | wwPDB |
| Map angles (°) | 90.0, 90.0, 90.0 | wwPDB |
| Pixel spacing (Å) | 0.59907, 0.59907, 0.59907 | Depositor |

Not For Manuscript Review

### 5 Model quality (i)

#### 5.1 Standard geometry (i)

The Z score for a bond length (or angle) is the number of standard deviations the observed value is removed from the expected value. A bond length (or angle) with $|Z| > 5$ is considered an outlier worth inspection. RMSZ is the root-mean-square of all Z scores of the bond lengths (or angles).

| Mol | Chain | Bond lengths | | Bond angles | |
|---|---|---|---|---|---|
| | | RMSZ | #\|Z\| >5 | RMSZ | #\|Z\| >5 |
| 1 | A | 0.68 | 0/3910 | 0.95 | 12/5303 (0.2%) |
| 1 | B | 0.69 | 0/3910 | 0.96 | 13/5303 (0.2%) |
| 1 | C | 0.69 | 0/3910 | 0.95 | 12/5303 (0.2%) |
| 1 | D | 0.68 | 0/3910 | 0.96 | 15/5303 (0.3%) |
| All | All | 0.69 | 0/15640 | 0.95 | 52/21212 (0.2%) |

Chiral center outliers are detected by calculating the chiral volume of a chiral center and verifying if the center is modelled as a planar moiety or with the opposite hand.A planarity outlier is detected by checking planarity of atoms in a peptide group, atoms in a mainchain group or atoms of a sidechain that are expected to be planar.

| Mol | Chain | #Chirality outliers | #Planarity outliers |
|---|---|---|---|
| 1 | A | 0 | 1 |
| 1 | B | 0 | 1 |
| 1 | C | 0 | 1 |
| 1 | D | 0 | 1 |
| All | All | 0 | 4 |

There are no bond length outliers.

All (52) bond angle outliers are listed below:

| Mol | Chain | Res | Type | Atoms | Z | Observed(°) | Ideal(°) |
|---|---|---|---|---|---|---|---|
| 1 | D | 459 | ARG | NE-CZ-NH2 | 8.66 | 124.63 | 120.30 |
| 1 | C | 459 | ARG | NE-CZ-NH2 | 8.18 | 124.39 | 120.30 |
| 1 | D | 475 | ARG | NE-CZ-NH2 | 7.80 | 124.20 | 120.30 |
| 1 | A | 459 | ARG | NE-CZ-NH2 | 7.44 | 124.02 | 120.30 |
| 1 | C | 475 | ARG | NE-CZ-NH2 | 7.43 | 124.01 | 120.30 |
| 1 | B | 475 | ARG | NE-CZ-NH2 | 7.08 | 123.84 | 120.30 |
| 1 | A | 349 | ARG | NE-CZ-NH2 | 7.07 | 123.83 | 120.30 |
| 1 | B | 459 | ARG | NE-CZ-NH2 | 7.01 | 123.81 | 120.30 |
| 1 | B | 413 | ARG | NE-CZ-NH2 | 6.84 | 123.72 | 120.30 |
| 1 | B | 241 | ARG | NE-CZ-NH2 | 6.81 | 123.71 | 120.30 |

*Continued on next page...*

Not For Manuscript Review

*Continued from previous page...*

| Mol | Chain | Res | Type | Atoms | Z | Observed(°) | Ideal(°) |
|---|---|---|---|---|---|---|---|
| 1 | D | 218 | ARG | NE-CZ-NH2 | 6.77 | 123.68 | 120.30 |
| 1 | A | 220 | ARG | NE-CZ-NH2 | 6.76 | 123.68 | 120.30 |
| 1 | B | 220 | ARG | NE-CZ-NH2 | 6.70 | 123.65 | 120.30 |
| 1 | A | 241 | ARG | NE-CZ-NH2 | 6.54 | 123.57 | 120.30 |
| 1 | B | 349 | ARG | NE-CZ-NH2 | 6.51 | 123.56 | 120.30 |
| 1 | C | 146 | ARG | NE-CZ-NH2 | 6.41 | 123.51 | 120.30 |
| 1 | C | 220 | ARG | NE-CZ-NH2 | 6.31 | 123.45 | 120.30 |
| 1 | C | 349 | ARG | NE-CZ-NH2 | 6.30 | 123.45 | 120.30 |
| 1 | C | 241 | ARG | NE-CZ-NH2 | 6.29 | 123.44 | 120.30 |
| 1 | A | 146 | ARG | NE-CZ-NH2 | 6.22 | 123.41 | 120.30 |
| 1 | B | 70 | TYR | CB-CG-CD2 | -6.22 | 117.27 | 121.00 |
| 1 | C | 413 | ARG | NE-CZ-NH2 | 6.18 | 123.39 | 120.30 |
| 1 | D | 146 | ARG | NE-CZ-NH2 | 6.07 | 123.34 | 120.30 |
| 1 | D | 241 | ARG | NE-CZ-NH2 | 6.03 | 123.32 | 120.30 |
| 1 | D | 70 | TYR | CB-CG-CD2 | -6.03 | 117.39 | 121.00 |
| 1 | D | 349 | ARG | NE-CZ-NH2 | 5.99 | 123.30 | 120.30 |
| 1 | B | 146 | ARG | NE-CZ-NH2 | 5.98 | 123.29 | 120.30 |
| 1 | D | 220 | ARG | NE-CZ-NH2 | 5.93 | 123.27 | 120.30 |
| 1 | A | 48 | TYR | CB-CG-CD1 | -5.91 | 117.45 | 121.00 |
| 1 | A | 402 | ARG | NE-CZ-NH2 | 5.89 | 123.25 | 120.30 |
| 1 | C | 127 | ARG | NE-CZ-NH2 | 5.87 | 123.24 | 120.30 |
| 1 | C | 70 | TYR | CB-CG-CD2 | -5.87 | 117.48 | 121.00 |
| 1 | D | 48 | TYR | CB-CG-CD1 | -5.84 | 117.49 | 121.00 |
| 1 | B | 48 | TYR | CB-CG-CD1 | -5.82 | 117.51 | 121.00 |
| 1 | D | 261 | ARG | NE-CZ-NH2 | 5.67 | 123.14 | 120.30 |
| 1 | B | 218 | ARG | NE-CZ-NH2 | 5.67 | 123.14 | 120.30 |
| 1 | C | 48 | TYR | CB-CG-CD1 | -5.59 | 117.64 | 121.00 |
| 1 | B | 127 | ARG | NE-CZ-NH2 | 5.59 | 123.09 | 120.30 |
| 1 | B | 261 | ARG | NE-CZ-NH2 | 5.55 | 123.08 | 120.30 |
| 1 | A | 70 | TYR | CB-CG-CD2 | -5.54 | 117.67 | 121.00 |
| 1 | A | 127 | ARG | NE-CZ-NH2 | 5.45 | 123.03 | 120.30 |
| 1 | D | 182 | ARG | NE-CZ-NH2 | 5.38 | 122.99 | 120.30 |
| 1 | A | 413 | ARG | NE-CZ-NH2 | 5.33 | 122.97 | 120.30 |
| 1 | C | 176 | ARG | NE-CZ-NH2 | 5.32 | 122.96 | 120.30 |
| 1 | A | 72 | ARG | NE-CZ-NH2 | 5.30 | 122.95 | 120.30 |
| 1 | B | 402 | ARG | NE-CZ-NH2 | 5.25 | 122.92 | 120.30 |
| 1 | C | 402 | ARG | NE-CZ-NH2 | 5.18 | 122.89 | 120.30 |
| 1 | D | 148 | ARG | NE-CZ-NH2 | 5.16 | 122.88 | 120.30 |
| 1 | A | 218 | ARG | NE-CZ-NH2 | 5.15 | 122.87 | 120.30 |
| 1 | D | 402 | ARG | NE-CZ-NH2 | 5.12 | 122.86 | 120.30 |
| 1 | D | 238 | ARG | NE-CZ-NH2 | 5.09 | 122.85 | 120.30 |
| 1 | D | 413 | ARG | NE-CZ-NH2 | 5.03 | 122.81 | 120.30 |

Not For Manuscript Review

There are no chirality outliers.

All (4) planarity outliers are listed below:

| Mol | Chain | Res | Type | Group |
|---|---|---|---|---|
| 1 | A | 68 | TYR | Sidechain |
| 1 | B | 68 | TYR | Sidechain |
| 1 | C | 68 | TYR | Sidechain |
| 1 | D | 68 | TYR | Sidechain |

#### 5.2 Too-close contacts (i)

In the following table, the Non-H and H(model) columns list the number of non-hydrogen atoms and hydrogen atoms in the chain respectively. The H(added) column lists the number of hydrogen atoms added and optimized by MolProbity. The Clashes column lists the number of clashes within the asymmetric unit, whereas Symm-Clashes lists symmetry-related clashes.

| Mol | Chain | Non-H | H(model) | H(added) | Clashes | Symm-Clashes |
|---|---|---|---|---|---|---|
| 1 | A | 3812 | 3772 | 3769 | 0 | 0 |
| 1 | B | 3812 | 3772 | 3769 | 0 | 0 |
| 1 | C | 3812 | 3772 | 3769 | 0 | 0 |
| 1 | D | 3812 | 3772 | 3769 | 1 | 0 |
| All | All | 15248 | 15088 | 15076 | 1 | 0 |

The all-atom clashscore is defined as the number of clashes found per 1000 atoms (including hydrogen atoms). The all-atom clashscore for this structure is 0.

All (1) close contacts within the same asymmetric unit are listed below, sorted by their clash magnitude.

| Atom-1 | Atom-2 | Interatomic distance (Å) | Clash overlap (Å) |
|---|---|---|---|
| 1:D:465:ILE:HD12 | 1:D:465:ILE:H | 1.83 | 0.42 |

There are no symmetry-related clashes.

#### 5.3 Torsion angles (i)

##### 5.3.1 Protein backbone (i)

In the following table, the Percentiles column shows the percent Ramachandran outliers of the chain as a percentile score with respect to all PDB entries followed by that with respect to all EM entries.

Not For Manuscript Review

The Analysed column shows the number of residues for which the backbone conformation was analysed, and the total number of residues.

| Mol | Chain | Analysed | Favoured | Allowed | Outliers | Percentiles | |
|---|---|---|---|---|---|---|---|
| 1 | A | 456/462 (99%) | 439 (96%) | 17 (4%) | 0 | 100 | 100 |
| 1 | B | 456/462 (99%) | 439 (96%) | 17 (4%) | 0 | 100 | 100 |
| 1 | C | 456/462 (99%) | 434 (95%) | 22 (5%) | 0 | 100 | 100 |
| 1 | D | 456/462 (99%) | 433 (95%) | 23 (5%) | 0 | 100 | 100 |
| All | All | 1824/1848 (99%) | 1745 (96%) | 79 (4%) | 0 | 100 | 100 |

There are no Ramachandran outliers to report.

##### 5.3.2 Protein sidechains ⓘ

In the following table, the Percentiles column shows the percent sidechain outliers of the chain as a percentile score with respect to all PDB entries followed by that with respect to all EM entries.

The Analysed column shows the number of residues for which the sidechain conformation was analysed, and the total number of residues.

| Mol | Chain | Analysed | Rotameric | Outliers | Percentiles | |
|---|---|---|---|---|---|---|
| 1 | A | 417/417 (100%) | 415 (100%) | 2 (0%) | 86 | 86 |
| 1 | B | 417/417 (100%) | 412 (99%) | 5 (1%) | 67 | 67 |
| 1 | C | 417/417 (100%) | 413 (99%) | 4 (1%) | 73 | 73 |
| 1 | D | 417/417 (100%) | 413 (99%) | 4 (1%) | 73 | 73 |
| All | All | 1668/1668 (100%) | 1653 (99%) | 15 (1%) | 74 | 75 |

All (15) residues with a non-rotameric sidechain are listed below:

| Mol | Chain | Res | Type |
|---|---|---|---|
| 1 | A | 93 | THR |
| 1 | A | 152 | CYS |
| 1 | B | 73 | MET |
| 1 | B | 84 | SER |
| 1 | B | 93 | THR |
| 1 | B | 152 | CYS |
| 1 | B | 477 | LEU |
| 1 | C | 93 | THR |
| 1 | C | 152 | CYS |
| 1 | C | 157 | ASP |

*Continued on next page...*

Not For Manuscript Review

*Continued from previous page...*

| Mol | Chain | Res | Type |
|---|---|---|---|
| 1 | C | 477 | LEU |
| 1 | D | 93 | THR |
| 1 | D | 152 | CYS |
| 1 | D | 157 | ASP |
| 1 | D | 449 | THR |

Sometimes sidechains can be flipped to improve hydrogen bonding and reduce clashes. There are no such sidechains identified.

##### 5.3.3 RNA (i)

There are no RNA molecules in this entry.

#### 5.4 Non-standard residues in protein, DNA, RNA chains (i)

There are no non-standard protein/DNA/RNA residues in this entry.

#### 5.5 Carbohydrates (i)

There are no oligosaccharides in this entry.

#### 5.6 Ligand geometry (i)

There are no ligands in this entry.

#### 5.7 Other polymers (i)

There are no such residues in this entry.

#### 5.8 Polymer linkage issues (i)

The following chains have linkage breaks:

| Mol | Chain | Number of breaks |
|---|---|---|
| 1 | C | 2 |
| 1 | D | 2 |
| 1 | B | 2 |
| 1 | A | 2 |

Not For Manuscript Review

All chain breaks are listed below:

| Model | Chain | Residue-1 | Atom-1 | Residue-2 | Atom-2 | Distance (Å) |
|---|---|---|---|---|---|---|
| 1 | C | 314:LEU | C | 325:ARG | N | 11.73 |
| 1 | D | 314:LEU | C | 325:ARG | N | 11.43 |
| 1 | B | 314:LEU | C | 325:ARG | N | 10.76 |
| 1 | A | 314:LEU | C | 325:ARG | N | 10.46 |
| 1 | A | 115:LYS | C | 126:ASN | N | 8.92 |
| 1 | D | 115:LYS | C | 126:ASN | N | 8.67 |
| 1 | B | 115:LYS | C | 126:ASN | N | 8.27 |
| 1 | C | 115:LYS | C | 126:ASN | N | 8.22 |

Not For Manuscript Review

### 6 Map visualisation (i)

This section contains visualisations of the EMDB entry D_1000287791. These allow visual inspection of the internal detail of the map and identification of artifacts.

Images derived from a raw map, generated by summing the deposited half-maps, are presented below the corresponding image components of the primary map to allow further visual inspection and comparison with those of the primary map.

#### 6.1 Orthogonal projections (i)

##### 6.1.1 Primary map

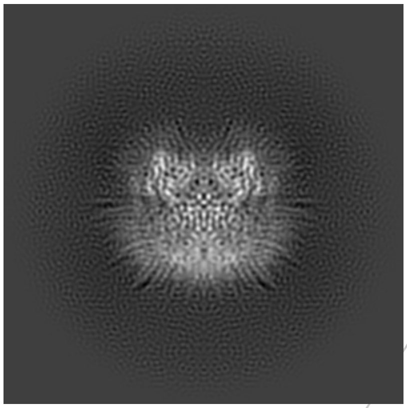

X

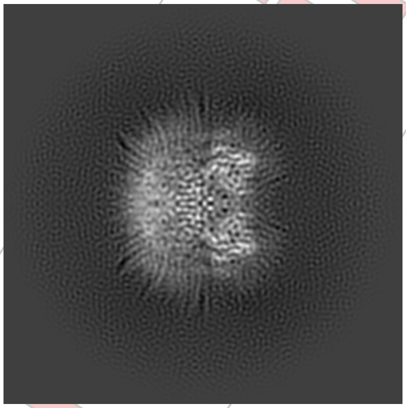

Y

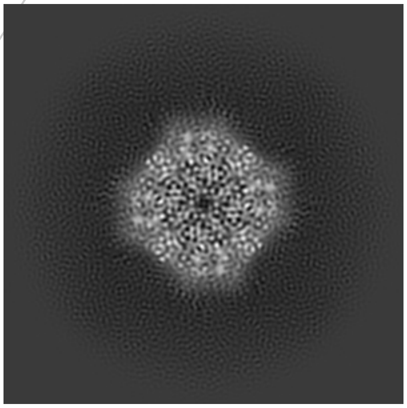

Z

##### 6.1.2 Raw map

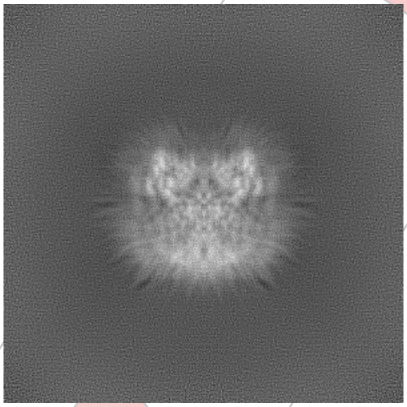

X

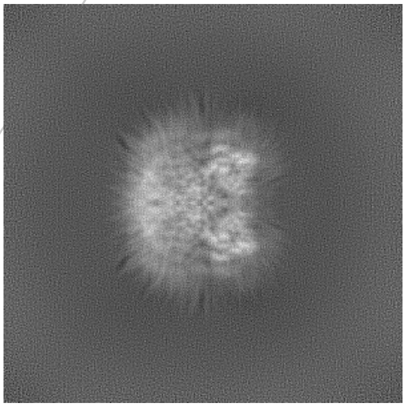

Y

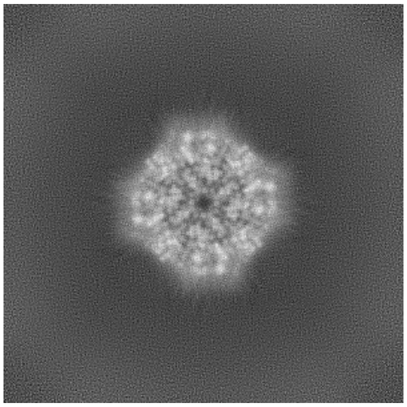

Z

The images above show the map projected in three orthogonal directions.

Not For Manuscript Review

#### 6.2 Central slices (i)

##### 6.2.1 Primary map

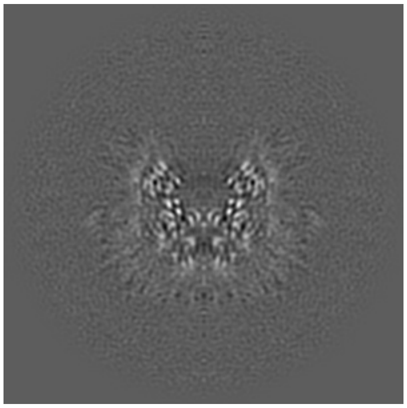
X Index: 216

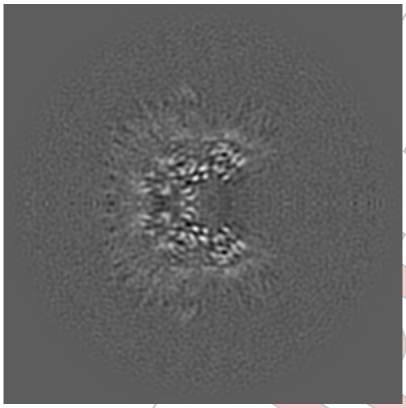
Y Index: 216

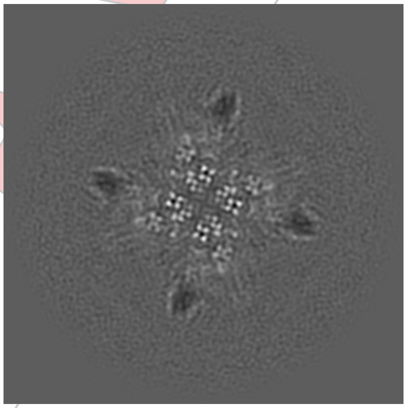
Z Index: 216

##### 6.2.2 Raw map

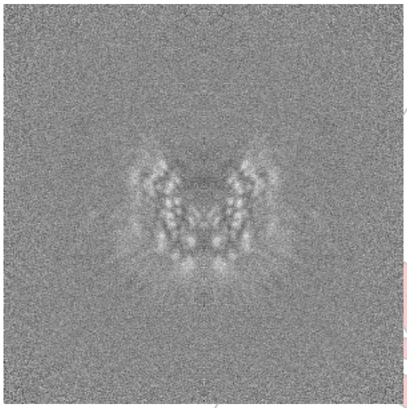
X Index: 216

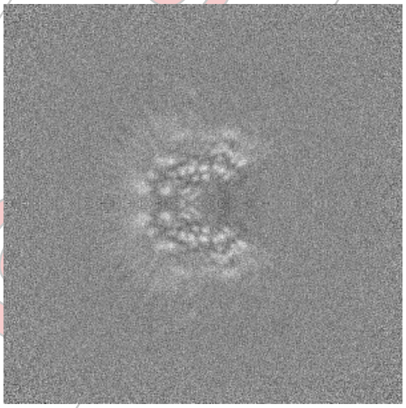
Y Index: 216

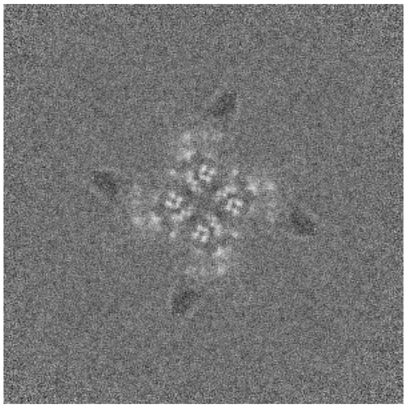
Z Index: 216

The images above show central slices of the map in three orthogonal directions.

Not For Manuscript Review

#### 6.3 Largest variance slices (i)

##### 6.3.1 Primary map

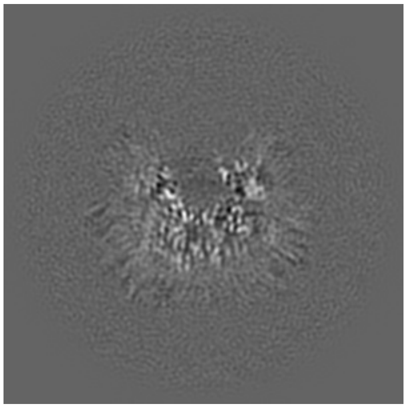
X Index: 210

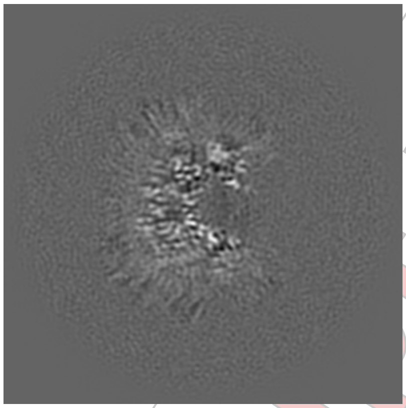
Y Index: 222

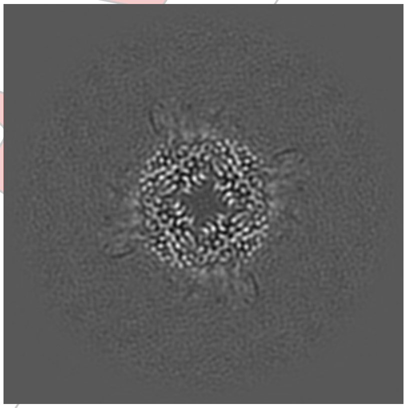
Z Index: 241

##### 6.3.2 Raw map

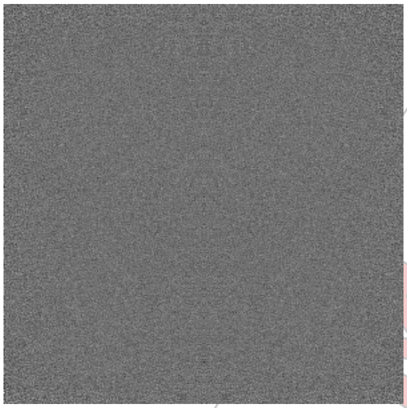
X Index: 0

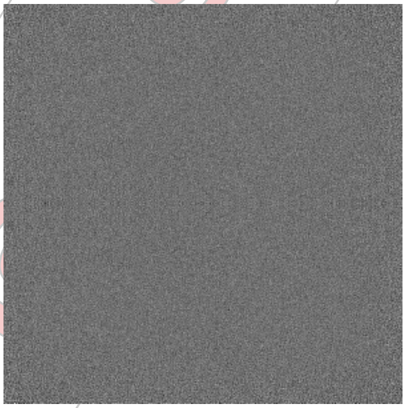
Y Index: 0

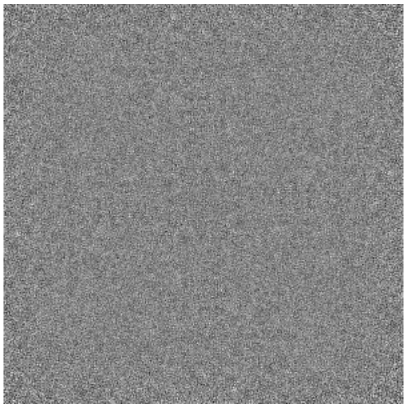
Z Index: 0

The images above show the largest variance slices of the map in three orthogonal directions.

Not For Manuscript Review

#### 6.4 Orthogonal standard-deviation projections (False-color) (i)

##### 6.4.1 Primary map

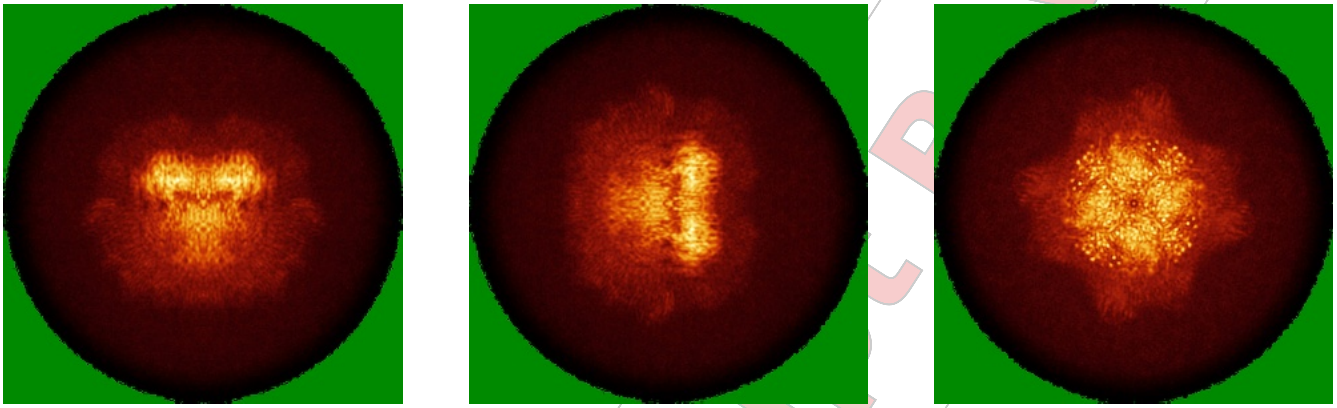

X Y Z

##### 6.4.2 Raw map

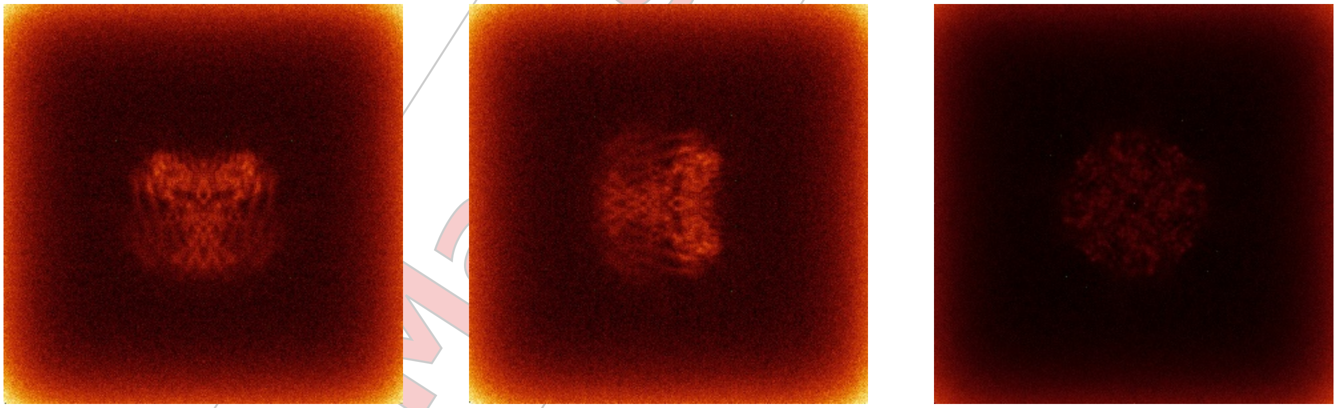

X Y Z

The images above show the map standard deviation projections with false color in three orthogonal directions. Minimum values are shown in green, max in blue, and dark to light orange shades represent small to large values respectively.

Not For Manuscript Review

Not For Manuscript Review

#### 6.5 Orthogonal surface views (i)

##### 6.5.1 Primary map

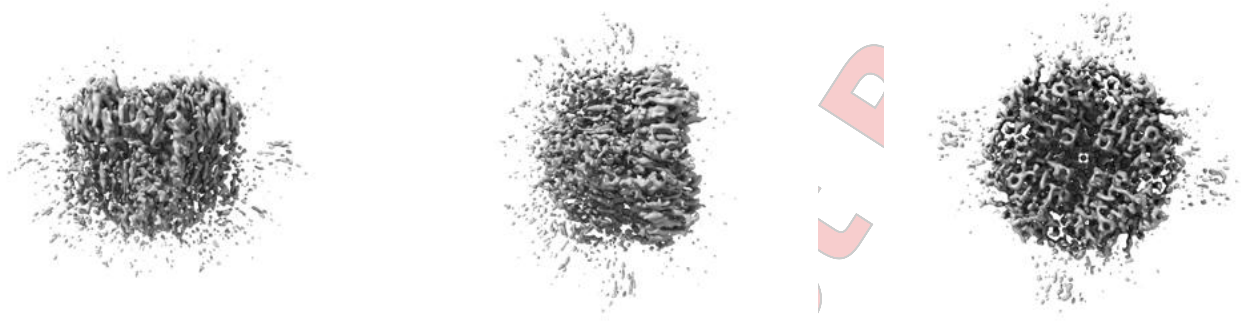

X Y Z

The images above show the 3D surface view of the map at the recommended contour level 0.0891. These images, in conjunction with the slice images, may facilitate assessment of whether an appropriate contour level has been provided.

##### 6.5.2 Raw map

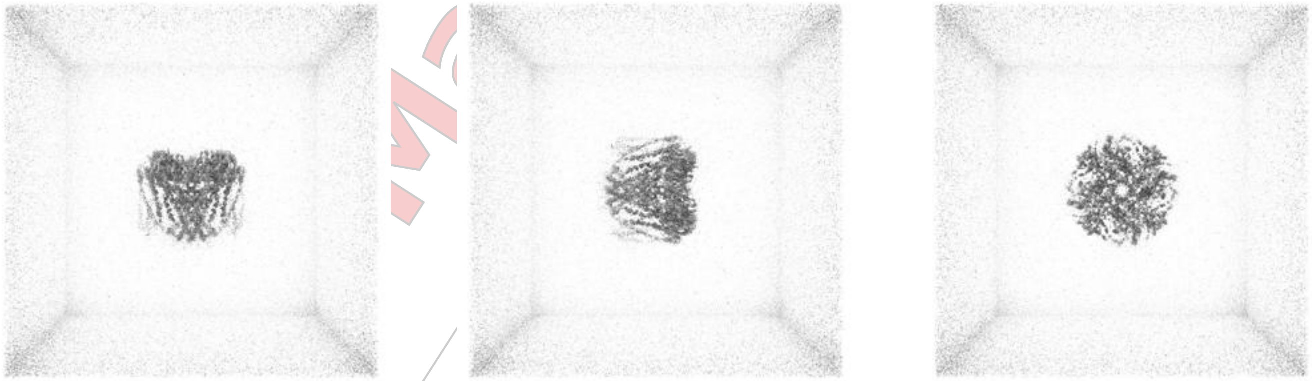

X Y Z

These images show the 3D surface of the raw map. The raw map's contour level was selected so that its surface encloses the same volume as the primary map does at its recommended contour level.

#### 6.6 Mask visualisation ⓘ

This section shows the 3D surface view of the primary map at 50% transparency overlaid with the specified mask at 0% transparency

A mask typically either:

- Encompasses the whole structure
- Separates out a domain, a functional unit, a monomer or an area of interest from a larger structure

##### 6.6.1 D_1000287791_em-mask-volume_P1.map.V2 ⓘ

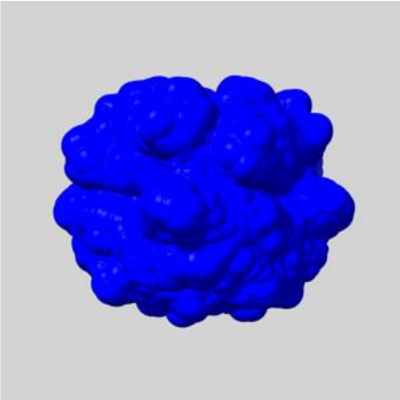

X

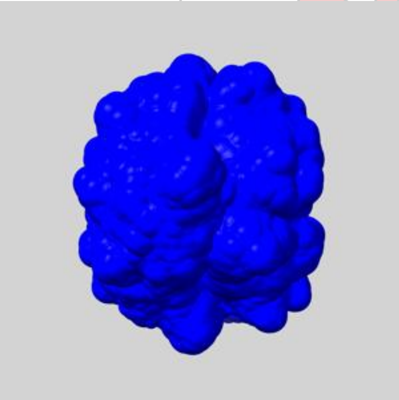

Y

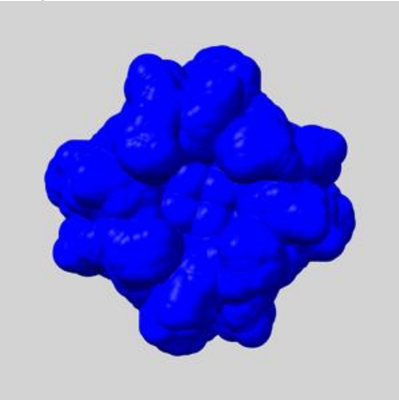

Z

Not For Manuscript Review

### 7 Map analysis (i)

This section contains the results of statistical analysis of the map.

#### 7.1 Map-value distribution (i)

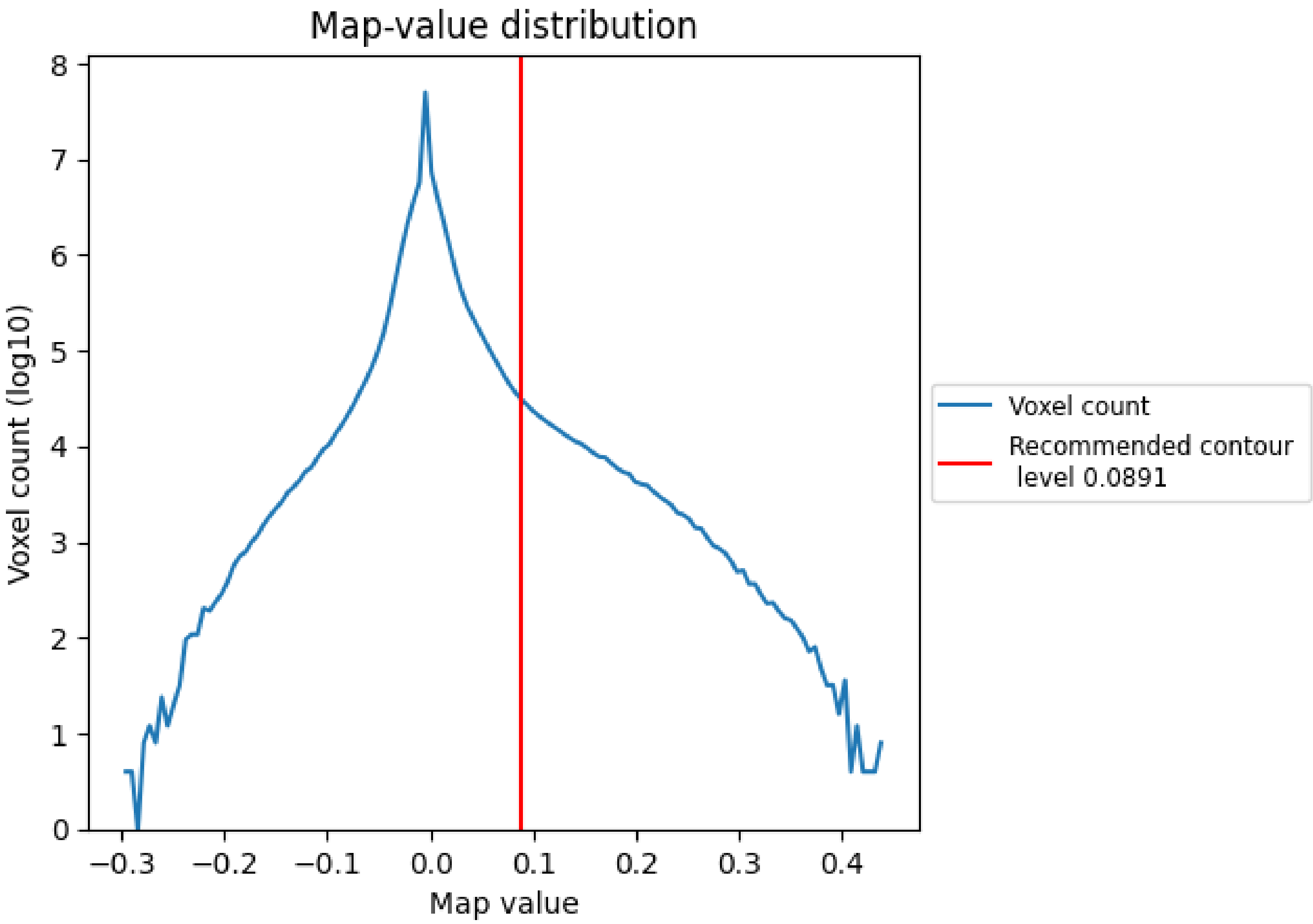


The map-value distribution is plotted in 128 intervals along the x-axis. The y-axis is logarithmic. A spike in this graph at zero usually indicates that the volume has been masked.

Not For Manuscript Review

#### 7.2 Volume estimate ⓘ

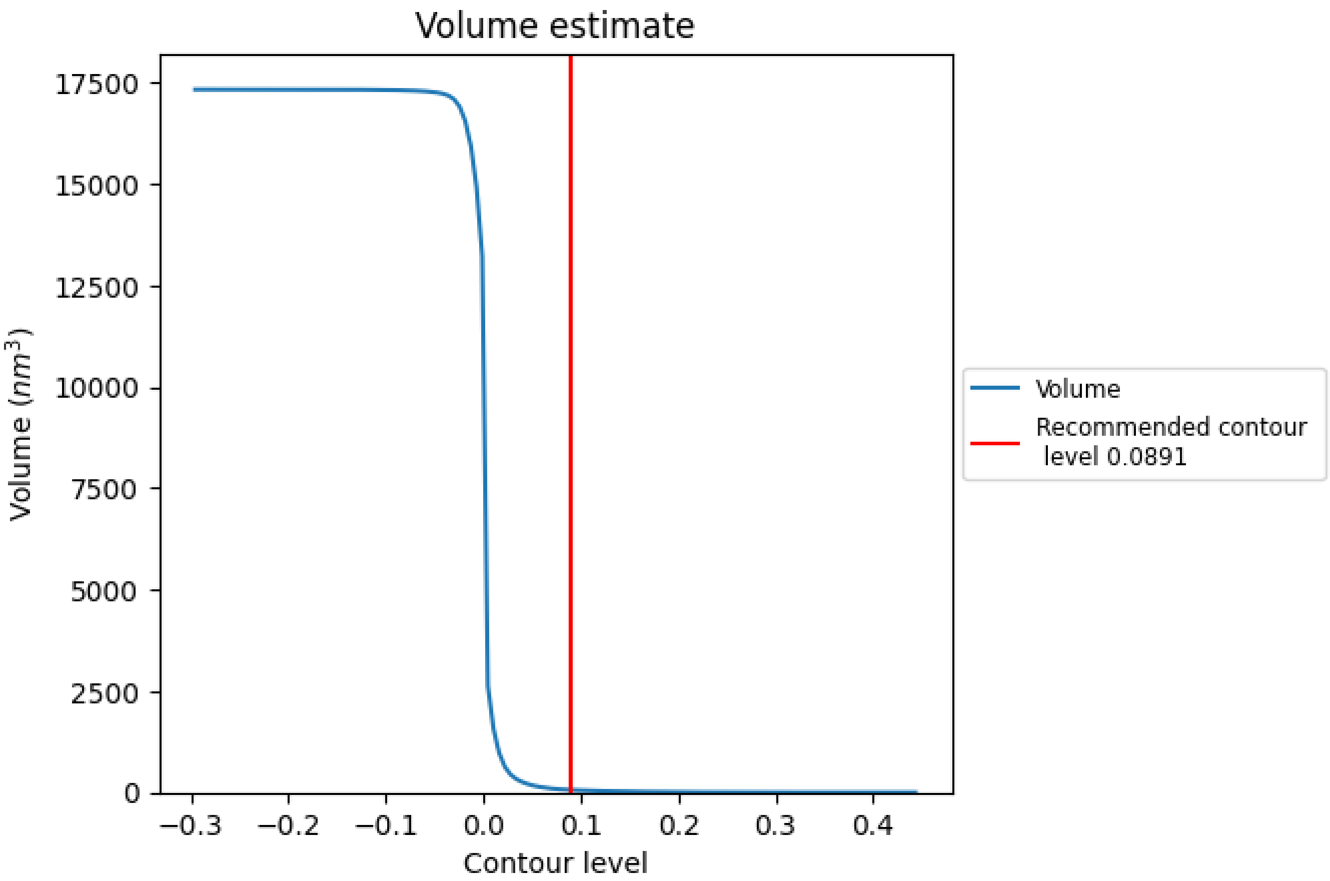


The volume at the recommended contour level is 63 $nm^3$; this corresponds to an approximate mass of 57 kDa.

The volume estimate graph shows how the enclosed volume varies with the contour level. The recommended contour level is shown as a vertical line and the intersection between the line and the curve gives the volume of the enclosed surface at the given level.

Not For Manuscript Review

#### 7.3 Rotationally averaged power spectrum (i)

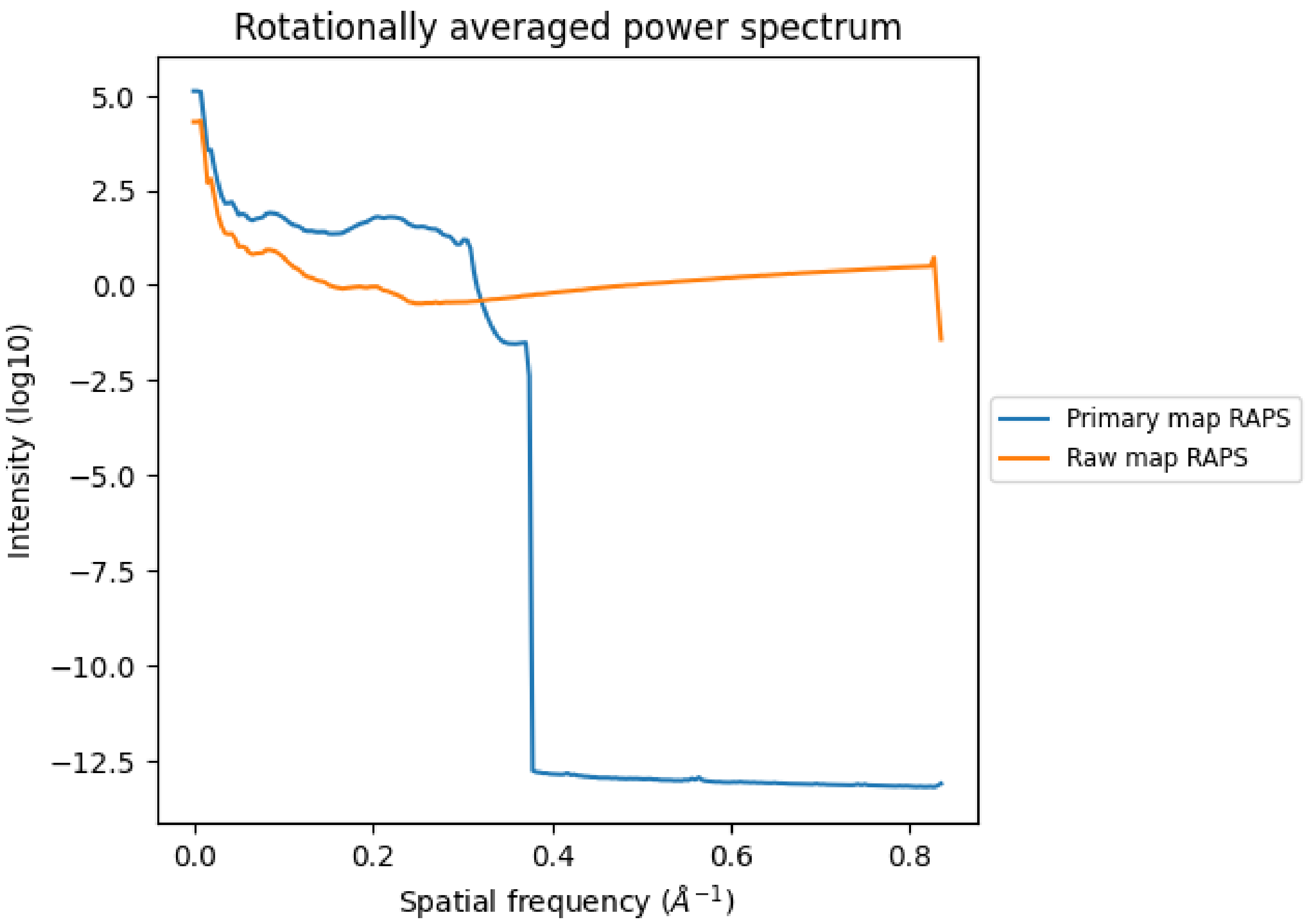


Not For Manuscript Review

### 8 Fourier-Shell correlation (i)

Fourier-Shell Correlation (FSC) is the most commonly used method to estimate the resolution of single-particle and subtomogram-averaged maps. The shape of the curve depends on the imposed symmetry, mask and whether or not the two 3D reconstructions used were processed from a common reference. The reported resolution is shown as a black line. A curve is displayed for the half-bit criterion in addition to lines showing the 0.143 gold standard cut-off and 0.5 cut-off.

#### 8.1 FSC (i)

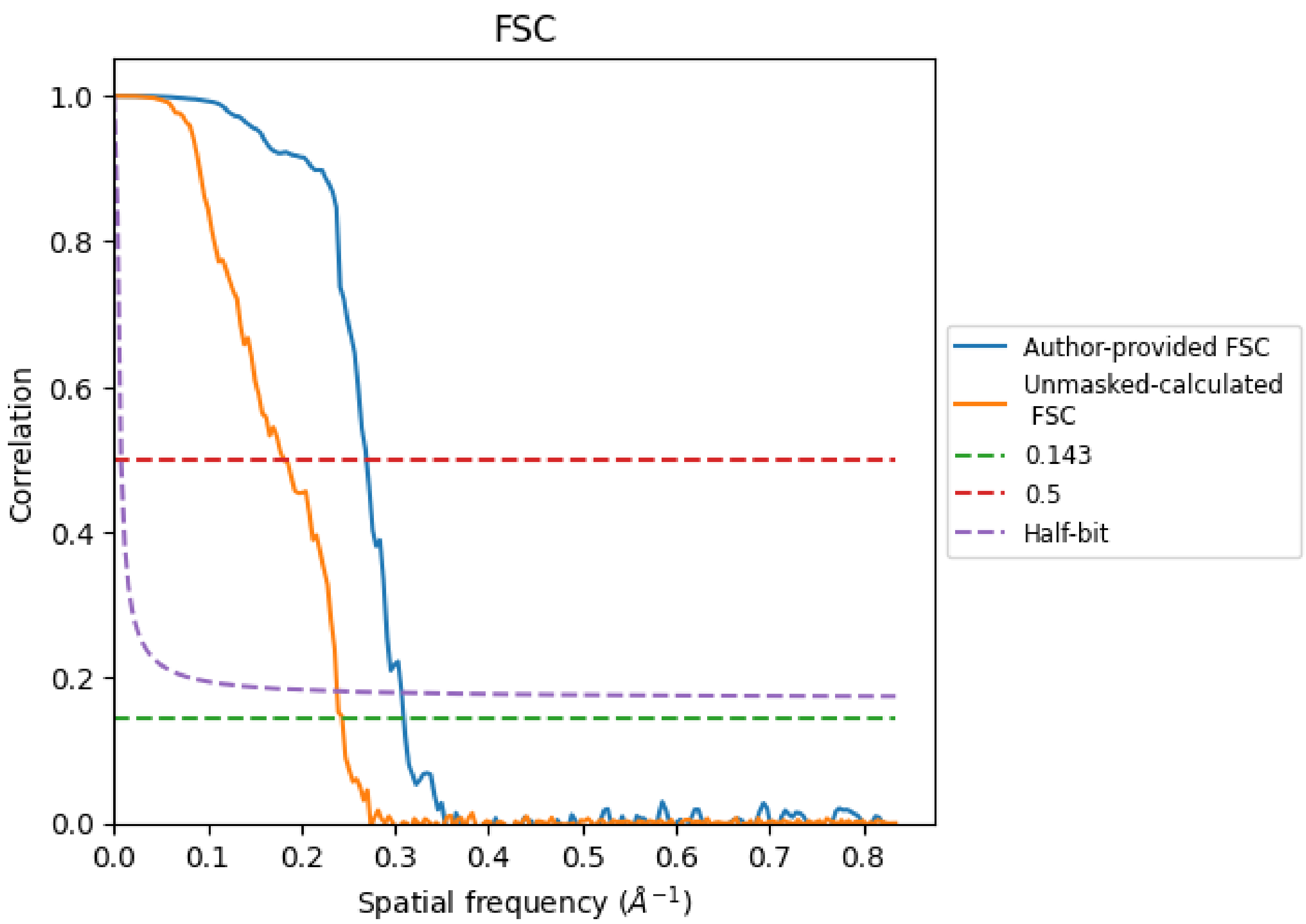


Not For Review

#### 8.2 Resolution estimates (i)

| Resolution estimate (Å) | Estimation criterion (FSC cut-off) | | |
|---|---|---|---|
| | 0.143 | 0.5 | Half-bit |
| Reported by author | - | - | - |
| Author-provided FSC curve | 3.23 | 3.71 | 3.26 |
| Unmasked-calculated* | 4.10 | 5.52 | 4.20 |

*Resolution estimate based on FSC curve calculated by comparison of deposited half-maps.

Not For Manuscript Review

### 9 Map-model fit (i)

This section contains information regarding the fit between EMDB map D_1000287791 and PDB model D_1000287791. Per-residue inclusion information can be found in section 3 on page 4.

#### 9.1 Map-model overlay (i)

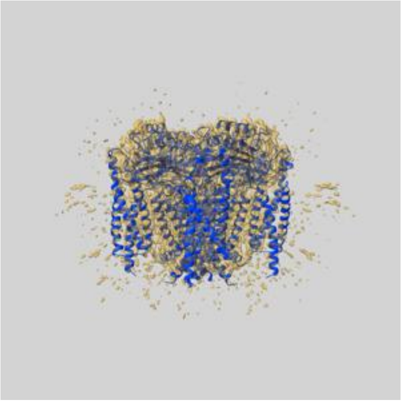

X

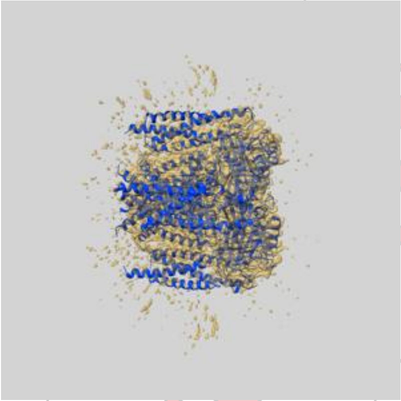

Y

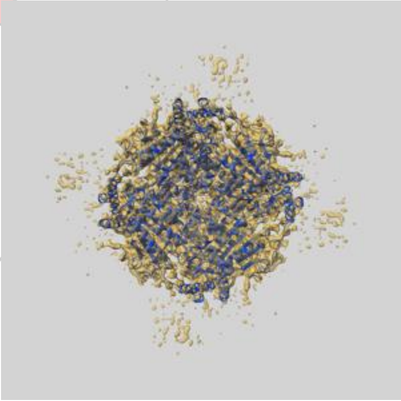

Z

The images above show the 3D surface view of the map at the recommended contour level 0.0891 at 50% transparency in yellow overlaid with a ribbon representation of the model coloured in blue. These images allow for the visual assessment of the quality of fit between the atomic model and the map.

Not For Manuscript Review

#### 9.2 Q-score mapped to coordinate model (i)

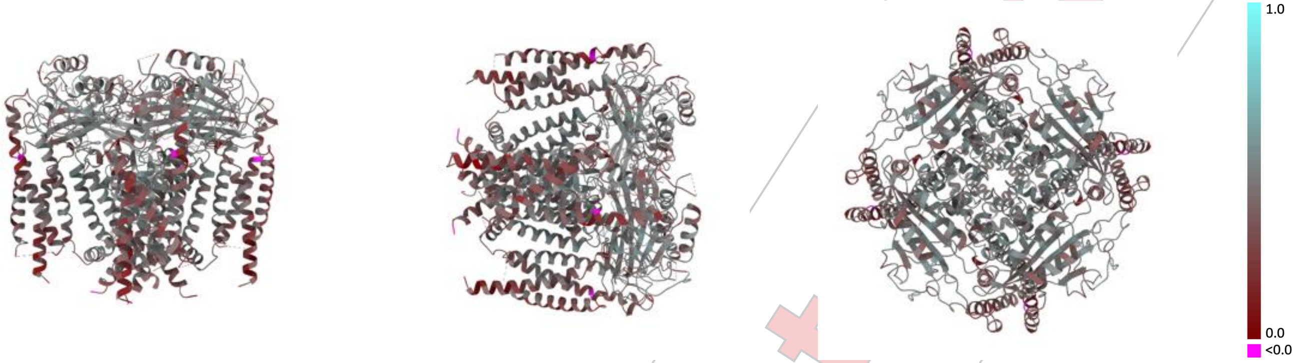


X Y Z

The images above show the model with each residue coloured according its Q-score. This shows their resolvability in the map with higher Q-score values reflecting better resolvability. Please note: Q-score is calculating the resolvability of atoms, and thus high values are only expected at resolutions at which atoms can be resolved. Low Q-score values may therefore be expected for many entries.

#### 9.3 Atom inclusion mapped to coordinate model (i)

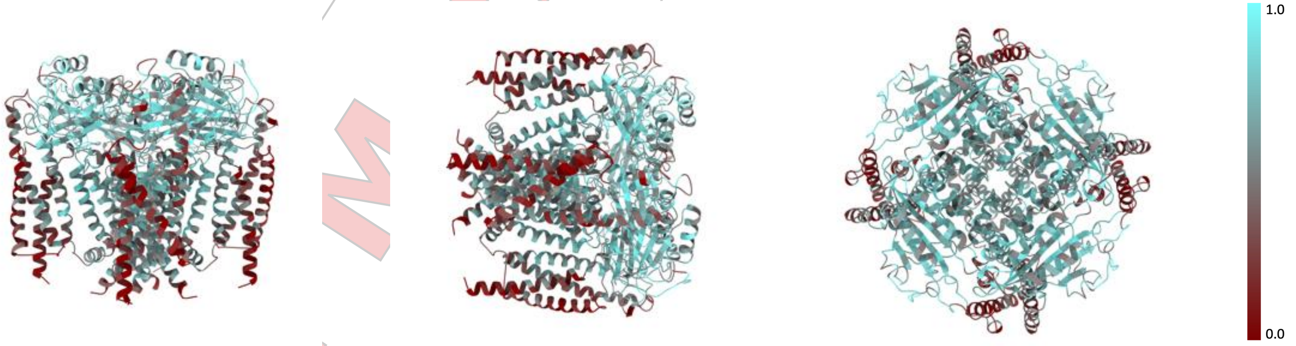


X Y Z

The images above show the model with each residue coloured according to its atom inclusion. This shows to what extent they are inside the map at the recommended contour level (0.0891).

Not For Manuscript Review

#### 9.4 Atom inclusion (i)

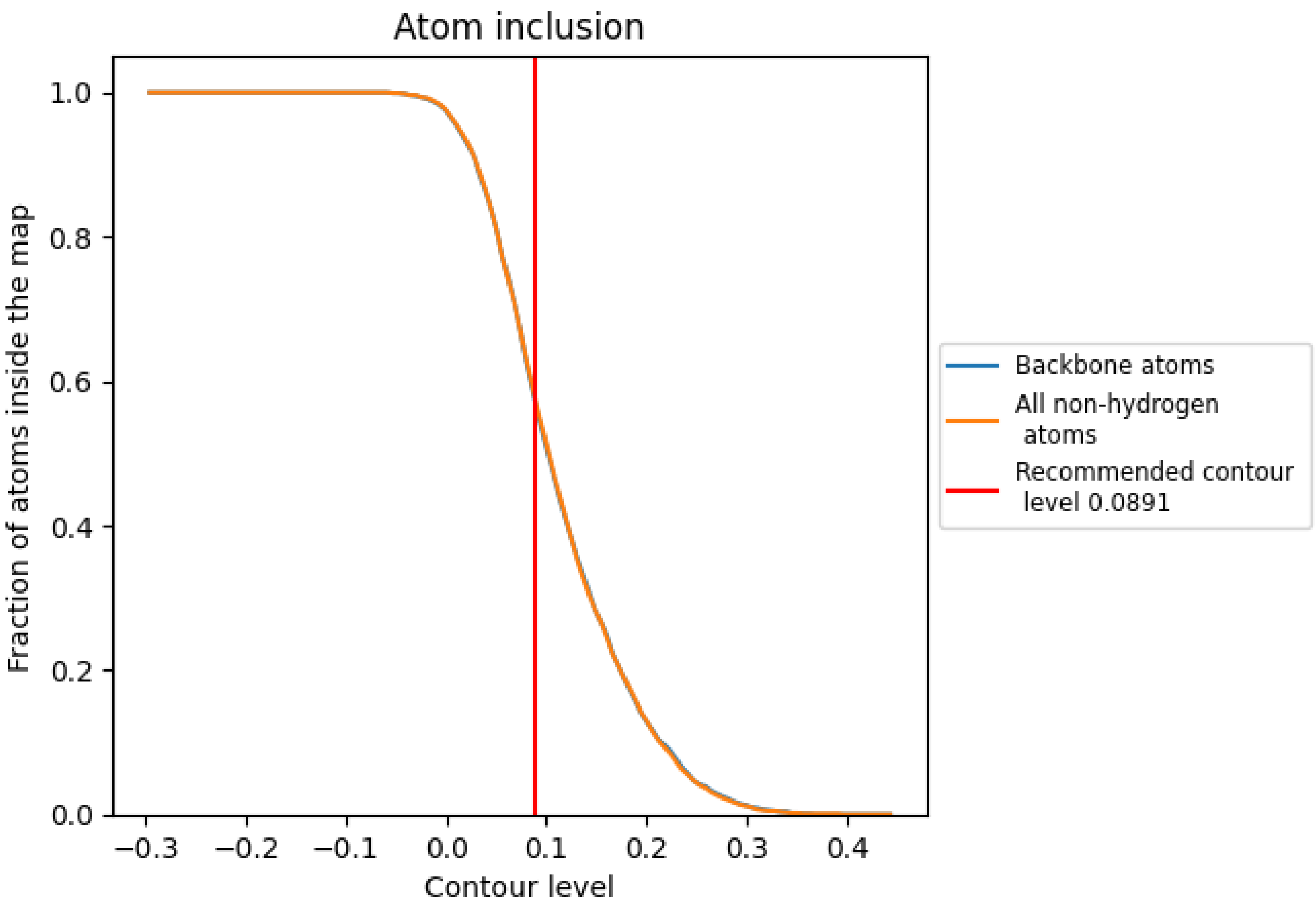


At the recommended contour level, 57% of all backbone atoms, 57% of all non-hydrogen atoms, are inside the map.

Not For Manuscript Review

#### 9.5 Map-model fit summary (i)

The table lists the average atom inclusion at the recommended contour level (0.0891) and Q-score for the entire model and for each chain.

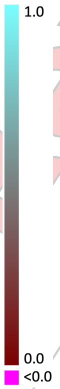


| Chain | Atom inclusion | Q-score |
|---|---|---|
| All | 0.5750 | 0.4250 |
| A | 0.5750 | 0.4260 |
| B | 0.5750 | 0.4250 |
| C | 0.5740 | 0.4260 |
| D | 0.5710 | 0.4250 |

Not For Manuscript Review
